# Study on *Dianthus superbus* L. against Acute Liver Injury via the Gut–Liver Axis — Regulation of Gut Microbiota and Tryptophan Metabolism to Activate AhR Signaling

**DOI:** 10.1101/2025.11.14.688543

**Authors:** Xiaolei Jiang, Yafeng Zhuang, Tiancheng Meng, Tianwei Meng, Jiali Liu, Hongyu Meng, Hong Chang

## Abstract

**Background:** Dianthus superbus L. (DS) treats liver diseases, but its ALI mechanism via gut microbiota/metabolites is unclear.

**Purpose:** Investigate DS effects on CCl₄-induced ALI and mechanisms.

**Methods:** DS components (HPLC-MS/MS) were identified. ALI rats received DS. Liver/colon injury, fibrosis (biochemistry, histology, IHC, IF, WB for α-SMA, Collagen I, Occludin, ZO-1), serum metabolomics, fecal 16S rRNA, and tryptophan-targeted metabolomics were assessed. Antibiotics and FMT from DS-treated donors explored microbiota roles.

**Results:** 37 components, including Kaempferol and Aurantiamide, were identified. DS attenuated ALI, reducing hepatic α-SMA/Collagen I while increasing AhR/Cyp1a1, and upregulating colonic Occludin/ZO-1. FMT replicated DS benefits. Untargeted metabolomics linked effects to Linoleic/Tryptophan metabolism. 16S rRNA showed increased beneficial microbiota (HT002, Lactobacillus, Romboutsia); FMT confirmed Lactobacillus enrichment. DS normalized tryptophan metabolites (4-Hydroxybenzoic acid, L-Kynurenine, Indole-3-carboxaldehyde).

**Conclusion:** DS ameliorates ALI gut-microbiota-dependently by modulating tryptophan metabolism, enhancing intestinal barrier function, activating AHR signaling, and suppressing fibrosis biomarkers, indicating its potential as a microbiota modulator for ALI.

**Importance:** Acute liver injury is a serious health threat with limited treatment options. This study explores how a traditional medicinal herb, *Dianthus superbus* L. (DS), protects the liver through an unexpected route: the gut. We discovered that DS works by rebalancing the community of gut bacteria, which in turn improves the intestinal barrier and fine-tunes the body’s tryptophan metabolism. This cascade of effects ultimately activates a key cellular pathway (AhR) that reduces liver inflammation and fibrosis. Our findings not only clarify how this traditional remedy works but also highlight the critical role of the gut-liver axis, positioning DS as a promising, microbiota-targeting therapeutic candidate for preventing and treating liver disease.

## 1. Introduction

Acute liver injury(ALI) refers to a clinical syndrome where the liver sustains damage from various causes within a short period, leading to abnormal liver function. Its etiology is diverse, and its clinical manifestations and severity vary considerably. In severe cases, it can progress to acute liver failure, which is life-threatening^1–3^. The pathophysiology of ALI is complex and diverse, involving the combined effects of multiple factors. Firstly, direct cytotoxicity plays a significant role in hepatocellular damage. For instance, certain drugs or chemicals can directly affect hepatocytes, leading to intracellular oxidative stress, mitochondrial dysfunction, and disruption of the cell membrane, ultimately resulting in cell necrosis or apoptosis^4^. Additionally, immune-mediated injury serves as a significant mechanism in ALI. During ALI, excessive activation of the immune system can trigger the release of inflammatory cytokines, further exacerbating the damage to hepatocytes ^5^. Metabolic dysregulation also plays a pivotal role in ALI. Under certain circumstances, disturbances in the metabolic functions of hepatocytes lead to energy metabolism disorders and lipid peroxidation, changes that can intensify hepatocyte injury and death ^6^. For instance, impairments in fatty acid β-oxidation may result in lipid accumulation and oxidative stress, thereby promoting hepatocyte apoptosis ^7^. Without timely intervention, ALI may rapidly progress, ultimately leading to severe consequences, including liver fibrosis and even cirrhosis^8^. Therefore, prompt identification of the etiology, interruption of damaging factors, and adoption of effective therapeutic measures are crucial for preventing disease progression.

Recent studies have confirmed a close association between gut microbiota and the development of ALI^9^.In recent years, there has been increasing attention on the gut microbiome-liver axis and its relationship with intestinal flora, as well as their pathological significance^10^. Emerging evidence suggests that individuals with ALI exhibit a dysregulated microbial structure, which can affect the diversity of intestinal flora and the balance of its derived metabolites^11^. For instance, an increase in *Bacteroidia* abundance indicates the potential occurrence of liver injury^12^. while an increase in the abundance of flora such as *Lactobacillus* and *Akkermansiaceae* predicts improvement in ALI^13, 14^. *Lactobacillus*, a type of bacteria widely found in nature ^15^, can colonize in the intestine, inhibit the growth of harmful bacteria, and maintain the balance of intestinal microecology ^16^. It can also improve the expression of intestinal tight junction proteins and enhance intestinal barrier function ^17^. Tryptophan is an important microbial metabolite ^18^, and tryptophan metabolites produced by intestinal microbes play a crucial role in the intestinal immune barrier. For example, metabolites such as indole-3-acrylic acid, indole-3-propionic acid, and indole-3-acetic acid can activate the aryl hydrocarbon receptor (AhR), improve the Th17/Treg balance, and alleviate intestinal inflammation and immune responses ^19^. These metabolites can also regulate the function of intestinal epithelial cells, enhance the integrity of the intestinal barrier, and prevent the invasion of pathogens ^20^.

An increasing body of research indicates that the interaction between gut microbiota and natural products has become a pivotal area of study for elucidating the mechanisms by which natural products exert protective effects against liver injury ^21^. Gut microbiota possesses the ability to metabolize natural products, thereby influencing their bioavailability and efficacy in anti-liver injury ^22^. Conversely, natural products have the potential to prevent and treat ALI by modulating gut microbiota ^23^. DS represents the dried aerial parts of *Dianthus superbus* L., a plant belonging to the Caryophyllaceae family. In Mongolian medicine, it is known as Bashaga-Gaoyou. “Bashaga” is one of the primary herbal formulas used in Mongolian medical clinics for the treatment of liver diseases and is widely employed in the management of such conditions ^24^. DS serves as a key component in clinically utilized prescriptions for liver ailments, specifically in Digeda-4 Decoction and Qinggan Jiuwei Powder. Digeda-4 Decoction exhibits significant efficacy in treating both chronic and acute liver dysfunction ^25^, while Qinggan Jiuwei Powder can alleviate the degree of liver fibrosis induced by CCl_4_ in rats, protect hepatocytes, and inhibit the Raf/MEK/ERK signaling pathway ^26^. DS is rich in bioactive components such as polysaccharides, flavonoids, and dianthramide, which exhibit excellent antioxidant, anti-inflammatory, and anti-hepatic injury effects ^27^. Previous studies have demonstrated that its polysaccharides can inhibit oxidative stress and inflammatory responses, promoting hepatocyte regeneration and repair ^28^. Dianthramide is capable of enhancing the liver’s antioxidant capacity and facilitating hepatocyte growth and division. These bioactive components may exert their anti-hepatic injury effects by influencing the gut microbiota ^29^. However, the impact of DS on the gut microbiota associated with ALI remains unknown. The impact of DS on the gut microbiota associated with ALI and its underlying mechanisms of action are currently poorly understood.

This study investigated the molecular mechanisms underlying the intervention of DS in ALI, with a focus on its regulatory role based on intestinal microbiota. The active components of DS were analyzed using UPLC-Q-TOF-MS technology. Based on a CCl_4_-induced animal model of ALI, the integration of pharmacodynamic evaluation, pathological analysis of liver and colon, and immunofluorescence detection confirmed that DS significantly improved liver tissue damage. Further combining untargeted metabolomics and 16SrRNA sequencing techniques, we delved deeper into the regulatory network of DS on ALI. Existing research has confirmed that abnormalities in tryptophan metabolism lead to the accumulation of its metabolites, which subsequently act as endogenous ligands to activate the AhR signal transduction pathway. Western blotting analysis revealed that DS significantly downregulated the expression of liver tissue fibrosis markers (α-SMA, Collagen) and key proteins in the AhR/Cyp1a1 signaling pathway, while upregulating the expression levels of intestinal tight junction proteins (ZO-1, Occludin). Through the construction of a pseudo-germ-free model and the implementation of fecal microbiota transplantation (FMT) experiments, it was confirmed that DS exerts hepatoprotective effects by reshaping the structure of intestinal flora. Mechanistic studies indicated that DS promotes intestinal epithelial repair and enhances intestinal barrier function by regulating specific functional flora (such as *Lactobacillus spp*.). This study elucidated the multi-target mechanism of DS in treating ALI from the perspective of the “gut-liver axis”, providing experimental evidence for the development of microbiome-based therapeutic strategies for liver diseases.

## 2. Materials and Methods

### 2.1 Materials

DS was purchased from Qiao Chinese Herbal Medicine Decoction Pieces Co., Ltd. (Anguo City); AST, ALT, AKP, γ-GT were acquired from Kete Biotech; acetonitrile and methanol (analytical grade) were sourced from Fisher (USA); formic acid (analytical grade) was supplied by CNW (Germany); silybin was obtained from MADUS GMBH; ampicillin, neomycin, metronidazole, and vancomycin were procured from Macklin. RIPA lysis buffer and protease inhibitor (Jinke Biotechnology); BCA protein concentration determination kit (Jinke Biotechnology); 5×loading buffer (Jinke Biotechnology); Colorful 180 broad-spectrum protein Marker (11-180KD) (Thermo Fisher), catalog number: #26617; ECL Plus ultrasensitive luminescent liquid (Biosharp), catalog number: BL520B; 10× TBST (Jinke Biotechnology); skim milk powder (Beyotime), catalog number: P0216-1500g; methanol (Sinopharm), catalog number: 80080418; 10× transfer buffer (Jinke Biotechnology); 5× MOPS/MES electrophoresis buffer (Jinke Biotechnology); MerckMillipore PVDF membrane 0.45µm (Millipore), catalog number: IPVH00010; MerckMillipore PVDF membrane 0.2µm (Millipore), catalog number: ISEQ00010.

### 2.2 DS Component Analysis

#### 2.2.1 Drug Extraction

2kg of DS was weighed and soaked in 70% ethanol at a solid-liquid ratio of 1:10 for 30 minutes. It was then transferred to a large round-bottom flask, added with 70% ethanol (maintaining the solid-liquid ratio of 1:10), and soaked for 1 hour. The flask was placed in an electric heating mantle and heated to boiling. The temperature was adjusted to maintain a distillation rate of 1 drop per second for an extraction duration of 2 hours. After cooling to room temperature, the mixture was filtered using a Buchner funnel. The residue was reheated and refluxed twice using the same method. The filtered liquids were combined, concentrated under reduced pressure to obtain an extract, and then lyophilized in a freeze dryer. The lyophilized extract was ground into powder, weighed, and stored for future use.

#### 2.2.2 Chromatographic and Mass Spectrometric Conditions for DS Components

The detection system employed was the Obtrap Exploris 120, utilizing a Waters Acquity UPLC BEH column (2.1*100, 1.7μm). The mobile phase A consisted of 0.1% formic acid in water, while mobile phase B was acetonitrile. The column temperature was maintained at 30℃, and the flow rate was set at 0.4ml/min. The elution gradient conditions for DS were as follows: 0-5min: 95%A, 6-10min: 90%A, 11-18min: 50%A, 19-30min: 10%A, and 31-35min: 95%A. The injection volume was 1.0μL.

For mass spectrometry, the ion source was a HESI source operating in positive and negative ion detection modes. The sheath gas pressure was set at 35Arb, with an auxiliary gas flow rate of 10Arb. The spray voltage was 3500V, and the ion transfer tube temperature was maintained at 320℃. The auxiliary gas temperature was 300℃. The scanning mode was Fullscan-ddMS2, with a full MS resolution of 60000 and a dd-MS2 resolution of 15000. The scanning range was from 100-1500, and the collision energy was set at 30%. All mass spectra were collected using the Xcalibur 3.0 software on the Obtrap Exploris 120 quadrupole Orbitrap high-resolution mass spectrometer.

#### 2.2.3 Data Processing

Baseline filtering, peak identification, integration, retention time correction, and peak alignment were performed using Progenesis QI. The molecular ion and fragment ion information of each component peak were analyzed using Xcalibur 4.0 software. By combining the accurate mass number of compounds, secondary cleavage fragment spectra, HMDB database (https://hmdb.ca/), and Pubchem database (https://pubchem.ncbi.nlm.nih.gov/), the main chemical components of DS were identified.

### 2.4 Animals

#### 2.4.1 Animal Grouping and Treatment

Sixty SPF-grade SD male rats (provided by Sibeifu (Beijing) Biotechnology Co., Ltd.) with a body weight of 180 ± 20 g were acclimated and fed for 7 days, The study included six groups (10 rats each): control (CON), model (MOD), three dose groups (DSH, DSM, DSL), and a silymarin (Sily) group.. Experimental animals were licensed under SCXK (Beijing 2019-0010). The housing conditions were maintained at a temperature of 22-26°C, humidity of approximately 33%, and a 12-hour light-dark cycle. The animal experiments were approved by the Medical Ethics Committee of Baotou Medical College, Inner Mongolia University of Science and Technology (Approval No.: Ethical Approval Document of Baotou Medical College: BYDLR [2023] No. 38). The experiment lasted for 14 days. The MOD, DS, and Sily groups underwent modeling twice a week (40% CCl4 + corn oil), with an initial injection of 5 mg/kg followed by subsequent injections of 3 mg/kg, obtaining an ALI model^30^. The CON and MOD groups were administered 0.1% carboxymethyl cellulose sodium solution by gavage daily. The Sily group received 50 mg/kg of silymarin dissolved in 0.1% carboxymethyl cellulose sodium solution. The DS groups received doses of 1.575, 3.15, and 6.3 g/kg(Based on clinical dosages of 9-15g), respectively, dissolved in 0.1% carboxymethyl cellulose sodium and administered by gavage.

#### 2.4.2 Sample Collection

On the 14th day of the experiment, after 12 hours of food and water deprivation, fresh fecal samples were collected from the rats using aseptic technique and the extrusion method. The samples were immediately placed in liquid nitrogen for rapid freezing and then transferred to a -80°C ultra-low temperature freezer for storage. Anesthesia was induced using a 3% pentobarbital sodium solution (30 mg/kg), and once the anesthesia took effect, blood was collected from the abdominal aorta. The blood samples were centrifuged at 4500×g for 15 minutes at 4°C to separate the serum, which was then aliquoted into sterile 250 μL EP tubes and stored at -80°C for future use.

The rats were euthanized by cervical dislocation, and the liver tissue was quickly removed. A 2 cm³ tissue block from the portal area was immediately fixed in 4% paraformaldehyde solution for subsequent histopathological, immunohistochemical, and immunofluorescence detection. The remaining liver tissue was aliquoted into EP tubes, rapidly frozen in liquid nitrogen, and stored at -80°C for molecular biology analyses such as Western blotting.

After collection, the colonic tissue was flushed with pre-cooled physiological saline at 4°C to remove luminal contents. Representative segments of the colon were fixed in 4% paraformaldehyde for histological analysis, while the remaining tissue was immediately frozen in liquid nitrogen and stored at -80°C for subsequent detection methods such as Western blot.

#### 2.4.3 Experiment on Germ-free Rats

In this experiment, 40 germ-free rats were divided into four groups: the control group (CON), the model pseudo-germ-free group (AMOD), the model pseudo-germ-free Dianthus superbus (ADS) group, and the model pseudo-germ-free Dianthus superbus fecal microbiota transplantation (AMF) group. For the germ-free rat experiment, an antibiotic mixture containing 0.2 g/L of ampicillin, neomycin, and metronidazole, as well as 0.1 g/L of vancomycin, was added to the drinking water throughout the entire experimental process.

For the FMT experiment, the recipient rats were first subjected to antibiotic mixture treatment for 10 days. After that, fresh feces were collected daily from the donor rats. The fecal samples were rapidly and uniformly mixed with sterile saline in a 1:9 ratio, centrifuged at 1000 x g, and the supernatant was collected and filtered through gauze. The recipient rats were then gavaged with a dose of 1 ml/kg. Modeling was performed after 10 days of antibiotic administration, using the same method as described above in “Animal Grouping”. Sample collection was conducted 14 days later, following the same method as outlined in “Sample Collection”.

#### 2.4.4 Biochemical Indices Determination and Pathological Study

Initially, liver and colon tissues fixed with 4% paraformaldehyde were embedded in paraffin. Hematoxylin and eosin were employed for pathological evaluation, while Masson staining was used to detect collagen protein in liver tissue, which was then observed under a microscope. According to the kit instructions, the levels of AST, ALT, AKP.

#### 2.4.5 Immunohistochemical Analysis

Immunohistochemistry was employed to detect the expression of α-SMA and Collagen-Ⅰ in liver tissue, as well as Occludin and ZO-1 in colonic tissue. Colon tissue was fixed using 4% paraformaldehyde, paraffin-embedded, and sectioned (2–3 μm). Tissue slides were washed with deionized water after dewaxing with xylene and rehydration with gradient ethanol. Antigenic repair procedures were performed using microwave and antigenic repair solutions. The slides were sealed with 10% normal goat serum for 30 min at room temperature. After that, they were incubated separately with the different primary antibodies at 4 °C overnight and then with the corresponding secondary antibodies at room temperature for 60 min. microscope. Six sections were selected from each group, and at least five non-overlapping and intact high-amplitude fields of view were randomly selected from each section. The average optical density and integrated optical density in each field of view were measured using Image-Pro Plus 6.0 software, and the mean values were calculated for statistical analysis.

#### 2.4.6 Immunofluorescence Analysis

Paraffin sections of rat liver tissue were dewaxed, underwent antigen retrieval, and were blocked with H_2_O_2_ and serum. The primary antibody, corresponding HRP-labeled secondary antibody, and fluorophore-labeled tyramine were added sequentially. After microwave retrieval treatment, the first round of primary and secondary antibodies were eluted, while the fluorophore-labeled tyramine remained attached to the target. When detecting the second and third targets, the previous steps were repeated for a new round of labeling and microwave retrieval treatment. The fourth primary antibody and 594-labeled fluorescent secondary antibody were added, followed by counterstaining of cell nuclei with DAPI. The slides were then covered with an antifading mounting medium. Finally, images were detected and collected using a slide scanner (Pannoramic, 3Dhistech, Hungary). DAPI emitted blue light, AhR emitted red light, and Cyp1a1 emitted green light. The number of positive cells for each index was analyzed using Image-Pro Plus 6.0 software.

#### 2.4.7 Western blot analysis

Approximately 0.1g of tissue sample was weighed and homogenized in 1mL of RIPA lysis buffer (with 10μL of PMSF added per 1mL of lysis buffer) using a tissue grinder. The homogenate was then lysed at 4℃ for 30 minutes, followed by centrifugation at 12,000 rpm for 10 minutes at 4℃. The supernatant was collected and subjected to BCA protein quantitation according to the kit instructions, and stored at - 80℃ for long-term preservation. Based on the BCA protein quantitation results, approximately 80μg of total protein was mixed with an appropriate amount of 5× protein loading buffer, denatured in boiling water for 10 minutes, and briefly centrifuged before loading. Freshly prepared electrophoresis buffer was used, and electrophoresis was performed at 140V until the loading dye just exited the resolving gel. The gel was then removed and equilibrated in 1× transfer buffer for 20 minutes. PVDF membranes and thin filter papers of appropriate size were cut, and the PVDF membrane was activated with methanol. Both the PVDF membrane and thin filter papers were equilibrated in the transfer buffer for 20 minutes. The stack was assembled in the order of negative electrode, thin filter paper, gel, PVDF membrane, thin filter paper, and positive electrode. Transfer was carried out at a constant current of 200mA for 1 hour. Blocking was performed using a 5% non-fat milk solution prepared in 1× TBST. Primary antibodies for AhR, Collagen 1, CYP1A1, Occludin, ZO-1, α-SMA (1:1000), and GAPDH (1:2000) were added according to the antibody instructions, and the membranes were incubated overnight at 4℃ on a shaking bed. The membranes were then washed three times with 1× TBST for 10 minutes each. Following this, the membranes were incubated with diluted secondary antibodies (goat anti-rabbit IgG-HRP, rabbit anti-goat IgG-HRP, goat anti-mouse IgG-HRP at 1:4000) for 1 hour at room temperature on a shaking bed. Finally, the membranes were washed three times with 1× TBST for 10 minutes each.

### 2.5 Untargeted metabolomics study of serum

#### 2.5.1 Serum Sample Preparation

Serum samples were retrieved from a -80°C freezer and thawed at 4°C. 200μl of rat serum was placed in a 1.5ml centrifuge tube and diluted with mass spectrometry-grade methanol at a 1:5 ratio, meaning 800μl of methanol was added. After vortexing for 30 seconds, the mixture was allowed to stand at 4°C for 2 hours. Subsequently, it was centrifuged at 13000rpm for 20 minutes, and the supernatant was filtered for membrane sampling. 50μl from all serum samples was pooled and mixed evenly to prepare a serum QC sample following the aforementioned sample preparation method.

#### 2.5.2 Chromatographic and Mass Spectrometric Conditions for Serum Samples

Chromatographic conditions were as follows: The UPLC-MS detection system was employed with an ACQUITY UPLC HSS T3 C18 column (2.1mm×100mm, 1.8μm, Waters). Mobile phase A consisted of acetonitrile (0.1% formic acid), while mobile phase B was water (0.1% formic acid). The column temperature was maintained at 30°C. Gradient elution conditions were set at 0-4min, 5%-40% A; 4-10min, 40%-99% A; 10-12min, 99% A. The injection volume was 5μL. Mass spectrometric conditions included an ESI source with positive and negative ion detection modes, a sheath gas pressure of 206.84 kPa, an auxiliary gas flow rate of 15.22 L/min, a spray voltage of 3.5 kV, an ion transfer tube temperature of 320°C, an auxiliary gas temperature of 350°C, and a column temperature of 40°C. The scanning mode was full MS/dd-MS2 with a full scan resolution of 70000, AGC target of 1e6, scan range of 100 to 1000 m/z, and a second-level scan (dd-MS2/dd-SIM) resolution of 17500, AGC target of 1e5, maximum IT of 50ms, and collision energies of 30, 60, and 90 eV, respectively.

#### 2.5.3 Data Processing

Raw data were analyzed using Progenesis QI software to obtain normalized unlabeled results for metabolic peaks in each sample. Samples were normalized based on the total ion intensity of each chromatogram. The data matrix, consisting of sample codes, RT-m/z pairs, and peak areas, was imported into EZinfo 3.0 for Waters software for principal component analysis (PCA), partial least squares discriminant analysis (PLS-DA), and orthogonal partial least squares discriminant analysis (OPLS-DA). Based on the OPLS-DA analysis results of the control and model groups, a variable importance in projection (VIP) list file was created to identify metabolic variables with VIP > 1 and an independent sample t-test p < 0.05 for further analysis. The structures of potential biomarkers for ALI were identified by combining public databases such as HMDB (http://www.hmdb.ca/), KEGG (http://www.genome.jp/kegg/), mbrole2 (http://csbg.cnb.csic.es/mbrole2/index.php), and relevant literature. The names of the identified biomarkers were imported into the Metaboanalyst 6.0 website (http://www.metaboanalyst.ca/) to analyze and determine the disrupted metabolic pathways. Metabolic pathways showing significant relevance to ALI were screened using an impact threshold criterion (> 0).

### 2.6 Targeted metabolomics study of fecal tryptophan

#### 2.6.1 Fecal Sample Preparation

Fecal samples were accurately weighed and mixed with 80% methanol in water. Steel balls were then added for grinding for 3 minutes, followed by extraction using a vortex mixer for 20 minutes. Afterward, the mixture was centrifuged at 12,000 rpm for 15 minutes at 4℃. The entire supernatant was transferred to a 1.5 mL eP tube and dried using a freeze concentrator. It was then reconstituted with 100 μL of the initial mobile phase, centrifuged again at 12,000 rpm for 15 minutes at 4℃, and the supernatant was collected for LC-MS/MS analysis.

#### 2.6.2 Chromatographic and Mass Spectrometric Conditions for Fecal Samples

In this study, an Agilent Poroshell 120 EC-C18 column (3.0 mm × 150 mm, 2.7 μm particle size) was used, operated at a column temperature of 40℃. The injection volume was set to 5 μL. Mobile phase A consisted of 5 mmol/L ammonium acetate in water (containing 0.01% formic acid), while mobile phase B was acetonitrile. Detection was performed using an electrospray ionization (ESI) source in both positive and negative ion modes. Nitrogen was used as the desolvation drying gas at a temperature of 325℃ and a flow rate of 8 L·min^-1. The capillary voltage was set to 3,500 V, the collision voltage to 135 V, and the nebulizer pressure to 35 psig. The scan range for MS1 was set from m/z 100 to 1,200, while the scan range for MS2 was from m/z 50 to 1,000. Mass spectrometry data were collected using Xcalibur 3.0 software on a UPLC-Q-Exactive quadrupole orbitrap high-resolution mass spectrometer.

#### 2.6.3 Data Processing

The raw metabolomics data were loaded and processed using Progenesis QI software. The data matrix was analyzed using EZinfor software, employing unsupervised principal component analysis (PCA) and supervised partial least squares-discriminant analysis (PLS-DA) to examine clustering and grouping. Differential metabolites were screened based on VIP > 1 in OPLS-DA combined with an independent samples t-test (with *p*<0.05). Cluster analysis of targeted differential metabolites was conducted using the MetaboAnalyst 6.0 platform. Changes in tryptophan-targeted metabolites across groups were analyzed using Graphpad 9.5 software.

### 2.7 Fecal 16S rDNA Analysis

#### 2.7.1 Fecal Sample Preparation

According to the requirements of 16S rRNA gene sequencing, Hangzhou LC-Bio Technologies Co., Ltd. extracted DNA from rat intestinal content samples using the E.Z.N.A® Stool DNA Kit. The extracted DNA was then subjected to agarose gel electrophoresis for quality assessment, and its quantity was determined using a UV spectrophotometer to ensure the accuracy and reliability of the results.

#### 2.7.2 Methods

The V3-V4 region of the prokaryotic small subunit (16S) rRNA gene was amplified using primers 341F (5’-CCTACGGGNGGCWGCAG-3’) and 805R (5’-GACTACHVGGGTATCTAATCC-3’). The PCR amplification was performed in a total volume of 25 μL reaction mixture containing 25 ng of template DNA, 12.5 μL of reaction buffer, 2.5 μL of each primer, and ddH2O to adjust the volume. The PCR products were then visualized using 2% agarose gel electrophoresis to confirm the presence of the desired DNA fragments. Ultra-pure water, instead of sample solution, was used during the entire DNA extraction process to eliminate the possibility of false-positive PCR results and served as a negative control. The PCR products were purified using AMPure XT beads (Beckman Coulter Genomics, Danvers, MA, USA) and quantified using Qubit (Invitrogen, USA). The amplified libraries were pooled for sequencing, and the size and quantity of the libraries were evaluated using an Agilent 2100 Bioanalyzer (Agilent, USA) and the Kapa Library Quantification Kit (Kapa Biosciences, Woburn, MA, USA), respectively. The libraries were sequenced on the NovaSeq PE250 platform.

#### 2.7.3 Data Processing

The gut microbiota data were analyzed using QIIME2 software. Alpha diversity analysis was performed using Graphpad 9.5, while beta diversity and species composition analysis were conducted on the LC-Bio Cloud Platform (https://www.omicstudio.cn/tool). Additionally, LEfSe analysis was carried out on the BioinCloud platform (https://www.bioincloud.tech/task-meta) with an LDA score threshold of 3.5.

### 2.8 Statistical Methods

This study employed SPSS 22.0 software for statistical analysis. Data were expressed in the format of X ± s. To compare differences between two groups, a T-test was utilized. A p-value less than 0.05 indicated that the study results were statistically significant, demonstrating significant differences between the groups. For data involving more than two groups, SPASS 22.0 software was used to perform a one-way analysis of variance (ANOVA). The results were represented as x ± s. Biochemical indicators, targeted metabolomics data, and untargeted metabolomics data from all rat groups were analyzed using Graphpad 9.5 software. Immunohistochemistry results, Masson staining results, and immunofluorescence findings were analyzed via Image-Pro Plus 6.0 software.

## 3. Results

### 3.1 Analysis of the Constituents of Total Extract from DS

In both positive and negative ion modes, 37 DS components were detected and identified. These primarily include nine phenolic acids: 4-Methoxy-benzeneacetic acid, 4-hydroxy-benzeneacetic acid, 3-Hydroxy-4-methoxybenzoic acid, Caffeic acid, Trans-p-coumaric acid, Hydroferulic acid, Salicylic acid, Scopoletin, and (E)-4-Methoxycinnamic acid. Six flavonoids were identified, namely Kaempferol, Quercetin-3-O-glucoside, Rutin, Orientin, Chrysoeriol-7-O-glucoside, and Diosmetin. Additionally, there are four terpenoids: Ursolic acid, Asiatic acid, Madecassic acid, Loliolide, and Vetol. Other identified compounds include phenylpropanoids, esters, amides, and more. Specific components are listed in Table 1.

**Tab 1.** Identification of DS components based on ultra-high-performance liquid chromatography coupled with mass spectrometry.

| No | Name | Formule | m/z | RT(min) | Adduct | Annot.<br>DeltaMass [ppm] | MS2 |
| --- | --- | --- | --- | --- | --- | --- | --- |
| 1 | (+)-Pinoresinol | C<sub>20</sub>H<sub>22</sub>O<sub>6</sub> | 358.39 | 7.26 | ESI+ | -2.47 | 137.9869、160.0748 |
| 2 | (E)-4-methoxycinnamic acid | C<sub>10</sub>H<sub>10</sub>O<sub>3</sub> | 178.185 | 15.3 | ESI+ | -2.47 | 133.1007、161.1320、105.0695 |
| 3 | 1-Monopalmitin | C<sub>19</sub>H<sub>38</sub>O<sub>4</sub> | 330.503 | 14.49 | ESI+ | -2.51 | 107.0852、125.0964、165.1635 |
| 4 | 2,6-Di-tert-butyl-p-benzoquinone | C<sub>14</sub>H<sub>20</sub>O<sub>2</sub> | 220.307 | 12.07 | ESI+ | -2.69 | 133.1008、161.1320、203.1424 |
| 5 | 4-Methoxy-benzeneacetic acid | C<sub>9</sub>H<sub>10</sub>O<sub>3</sub> | 166.17 | 31.86 | ESI+ | -1.77 | 167.01 |
| 6 | Aurantiamide acetate | C<sub>27</sub>H<sub>28</sub>N<sub>2</sub>O<sub>4</sub> | 444.522 | 12.98 | ESI+ | -3.81 | 105.0332、194.1170、224.1063 |
| 7 | Aurantiamide | C<sub>25</sub>H<sub>26</sub>N<sub>2</sub>O<sub>3</sub> | 402.486 | 12.98 | ESI+ | -2.78 | 117.0695、152.1065、385.1898 |
| 8 | Benzyl benzoate | C<sub>14</sub>H<sub>12</sub>O<sub>2</sub> | 212.244 | 8.06 | ESI+ | -2.75 | 153.0979 |
| 9 | Dianthoside | C<sub>12</sub>H<sub>16</sub>O<sub>8</sub> | 288.25 | 18.55 | ESI+ | -3.17 | 139.1114 |
| 10 | Dibutyl phthalate | C<sub>16</sub>H<sub>22</sub>O<sub>4</sub> | 278.34 | 14.68 | ESI+ | -3.02 | 121.1007、151.1408、167.1064 |
| 11 | Eugenol | C<sub>10</sub>H<sub>12</sub>O<sub>2</sub> | 164.201 | 6.69 | ESI+ | -1.88 | 109.1009、137.0957、165.0904 |
| 12 | Kaempferol | C<sub>15</sub>H<sub>10</sub>O<sub>6</sub> | 286.236 | 12.97 | ESI+ | -1.89 | 121.0280、213.1632 |
| 13 | Lariciresinol | C<sub>20</sub>H<sub>24</sub>O<sub>6</sub> | 360.4 | 6.59 | ESI+ | -3.01 | 361.1734 |
| 14 | Loliolide | C<sub>11</sub>H<sub>16</sub>O<sub>3</sub> | 196.24 | 8.94 | ESI+ | -2.13 | 133.1007、161.0956、179.1060 |
| 15 | Orientin | C<sub>21</sub>H<sub>20</sub>O<sub>11</sub> | 448.377 | 3.73 | ESI+ | -2.77 | 271.0774、449.1931 |
| 16 | 2-Phenylethanol | C<sub>8</sub>H<sub>10</sub>O | 122.16 | 0.86 | ESI+ | -2.48 | 122.9242 |
| 17 | Phenylacetaldehyde | C<sub>8</sub>H<sub>8</sub>O | 120.148 | 2.22 | ESI+ | -2.12 | 103.0538、121.0959 |
| 18 | Scopoletin | C<sub>10</sub>H<sub>8</sub>O<sub>4</sub> | 192.168 | 8.85 | ESI+ | -1.93 | 133.1007、137.0956、193.1581 |
| 19 | Sorbic Acid | C<sub>6</sub>H<sub>8</sub>O<sub>2</sub> | 112.127 | 3.8 | ESI+ | -1.96 | 57.0334、72.0805、95.0488 |
| 20 | Trans-p-coumaric acid | C<sub>9</sub>H<sub>8</sub>O<sub>3</sub> | 164.158 | 6.58 | ESI+ | -2.94 | 119.0852、147.0799、165.0904 |
| 21 | Ursolic acid | C<sub>30</sub>H<sub>48</sub>O<sub>3</sub> | 456.7 | 11.09 | ESI+ | -2.69 | 95.0852、107.0851、163.1482 |
| 23 | 3-Hydroxy-4-methoxybenzoic acid | C<sub>8</sub>H<sub>8</sub>O<sub>4</sub> | 168.147 | 11.29 | ESI- | -2.69 | 125.0968、169.0866 |
| 24 | 4-hydroxy-benzeneacetic acid | C<sub>8</sub>H<sub>8</sub>O<sub>3</sub> | 152.147 | 6.69 | ESI- | -2.5 | 90.9234、91.9312、151.0610 |
| 25 | Asiatic acid | C<sub>30</sub>H<sub>48</sub>O<sub>5</sub> | 488.699 | 12.25 | ESI- | -2.44 | 487.3 |
| 26 | Caffeic acid | C<sub>9</sub>H<sub>8</sub>O<sub>4</sub> | 215.245 | 6.63 | ESI- | -2.25 | 135.0448、179.0708 |
| 27 | Chrysoeriol-7-O-glucoside | C<sub>22</sub>H<sub>22</sub>O<sub>11</sub> | 462.404 | 8.79 | ESI- | -2.17 | 298.0473、461.1071 |
| 28 | Diosmetin | C<sub>16</sub>H<sub>12</sub>O<sub>6</sub> | 300.263 | 19.9 | ESI- | -2.37 | 299.2582 |
| 29 | Emodin | C<sub>15</sub>H<sub>10</sub>O<sub>5</sub> | 270.237 | 16.58 | ESI- | -2.33 | 269.2115 |
| 30 | Hydroferulic acid | C<sub>10</sub>H<sub>12</sub>O<sub>4</sub> | 196.2 | 8.97 | ESI- | -2.32 | 119.9257、195.1018 |
| 31 | Madecassic acid | C<sub>30</sub>H<sub>48</sub>O<sub>6</sub> | 504.698 | 6.84 | ESI- | -1.94 | 503.1757 |
| 32 | Palmitic acid | C<sub>16</sub>H<sub>32</sub>O<sub>2</sub> | 256.424 | 12.73 | ESI- | -2.87 | 238.0504、256.0609 |
| 33 | Quercetin-3-O-glucoside | C<sub>21</sub>H<sub>20</sub>O<sub>12</sub> | 464.376 | 17.48 | ESI- | -2.05 | 97.1008、465.3347 |
| 34 | Rutin | C<sub>27</sub>H<sub>30</sub>O<sub>16</sub> | 610.518 | 19.1 | ESI- | -2.05 | 610.2717、611.2744 |
| 35 | Salicylic acid | C<sub>7</sub>H<sub>6</sub>O<sub>3</sub> | 138.121 | 4.75 | ESI- | -2.9 | 93.03、137.02 |
| 36 | Vetol | C<sub>6</sub>H<sub>6</sub>O<sub>3</sub> | 126.11 | 9.35 | ESI- | -2.35 | 125.0969 |
| 37 | Succinic acid | C<sub>4</sub>H<sub>6</sub>O<sub>4</sub> | 118.088 | 11.08 | ESI- | -2.84 | 73.0293、99.9256、116.9281 |

### 3.2 DS increased liver function indices in ALI rats

To evaluate the therapeutic effect of DS on CCL_4_-induced ALI rats, this study determined liver function indicators in the liver tissue of rats in each group (Fig 1 A). Rats in the MOD group showed significantly elevated levels of AST, ALT, and AKP (*p*<0.05). After DS treatment, the levels of ALT and AKP in ALI rats were significantly reduced (*p*<0.05). Based on this, it is inferred that DS improves liver function in ALI rats to some extent.

**Fig 1.**
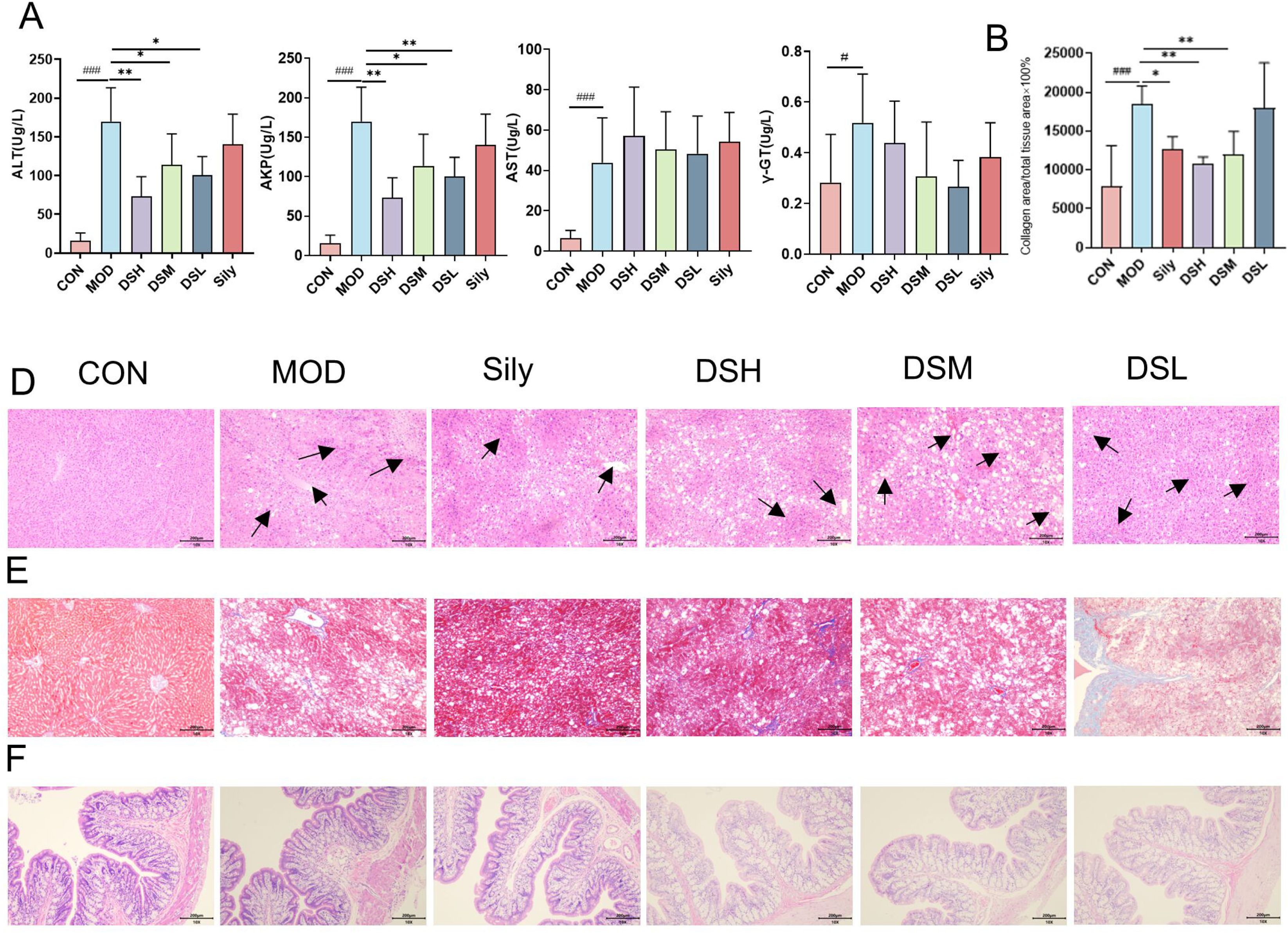
Biochemical and pathological effects of DS on ALI rats A: Biochemical indicators of liver tissue; B: Biochemical indicators of serum; C: Quantitative analysis of Masson staining in liver tissue; D: HE staining of liver tissue; E: Masson staining of liver tissue; F: HE staining of colonic tissue Compare with MOD:\*\*\**p*<0.001,\*\**p*<0.01,\**p*<0.05; Compare with CON:###*p*< 0.001,##*p*<0.01,#*p*<0.05

### 3.3 DS reduces liver and colonic tissue lesions in ALI rats

To evaluate the therapeutic effect of DS on ALI rats, this study conducted a detailed analysis of the liver tissues from each group of rats. In the HE-stained liver sections of CON rats (Fig 1 D), the hepatic cell structure was observed to be intact, with no evidence of degenerative or necrotic cells, and the lobular structure of the liver was clearly defined. Compared to CON, the MOD rats exhibited disruption of the lobular architecture, formation of pseudolobules, accompanied by substantial fatty degeneration and necrotic fibrous septa formation. Diffuse inflammatory cell infiltration was visible within the stroma. In the DS-treated rats, most of the hepatocytes remained intact, with only minimal fatty degeneration and inflammatory cells observed, particularly with the most pronounced effect in the DSM group. After processing the liver tissues from each group with Masson staining (Fig 1 E), collagen fibers appeared blue, muscle fibers appeared red, and cell nuclei appeared blue-black. The liver tissue sections from CON rats showed normal staining, whereas MOD rats presented with extensive blue areas, indicating prominent fibrotic symptoms. Following DS treatment, the blue areas in MOD rats were reduced, especially in the high and medium DS treatment groups, which demonstrated more significant effects (*p*<0.05) (Fig 1 B). Furthermore, intestinal barrier dysfunction is a common indicator of ALI. Therefore, this study further assessed the impact of DS on intestinal barrier function and performed HE staining on colon tissues. According to the staining results (Fig 1 F), DS treatment significantly improved the loss of goblet cells induced by ALI and contributed to restoring the homeostasis of the intestinal mucosa.

### 3.4 DS ameliorates ALI by modulating liver fibrotic proteins and intestinal barrier function proteins in the colon

Based on pathological findings and biochemical indicator analysis, we hypothesized that DSM would exhibit the most pronounced effects in improving ALI. Thus, we focused our subsequent investigations on elucidating the role of DSM in ALI. The analysis of α-SMA and Collagen I mRNA expression in liver tissue is crucial for assessing the severity of ALI. Accordingly, this study quantitatively analyzed the expression levels of α-SMA and Collagen I in rat liver tissue (Fig 3 A, B). Compared to CON rats, MOD rats demonstrated significantly increased expression of α-SMA and Collagen I in their liver tissue (*p* < 0.05). However, upon DSH treatment, there was a noticeable decrease in the expression levels of α-SMA and Collagen I in the liver tissue of ALI rats (*p* < 0.05).

To explore whether CCl_4_-induced ALI affects the integrity of the intestinal barrier, we analyzed the mRNA expression levels of ZO-1 and Occludin in rat colonic tissue (Fig 3 C, D). The results revealed a significant reduction in the mRNA expression levels of ZO-1 and Occludin in the colonic tissue of MOD rats compared to CON rats (*p* < 0.05). Following DSM treatment, the expression levels of ZO-1 and Occludin were significantly elevated (*p* < 0.05), thereby attenuating the increase in intestinal permeability induced by ALI in rats (*p* < 0.05).

**Fig 2.**
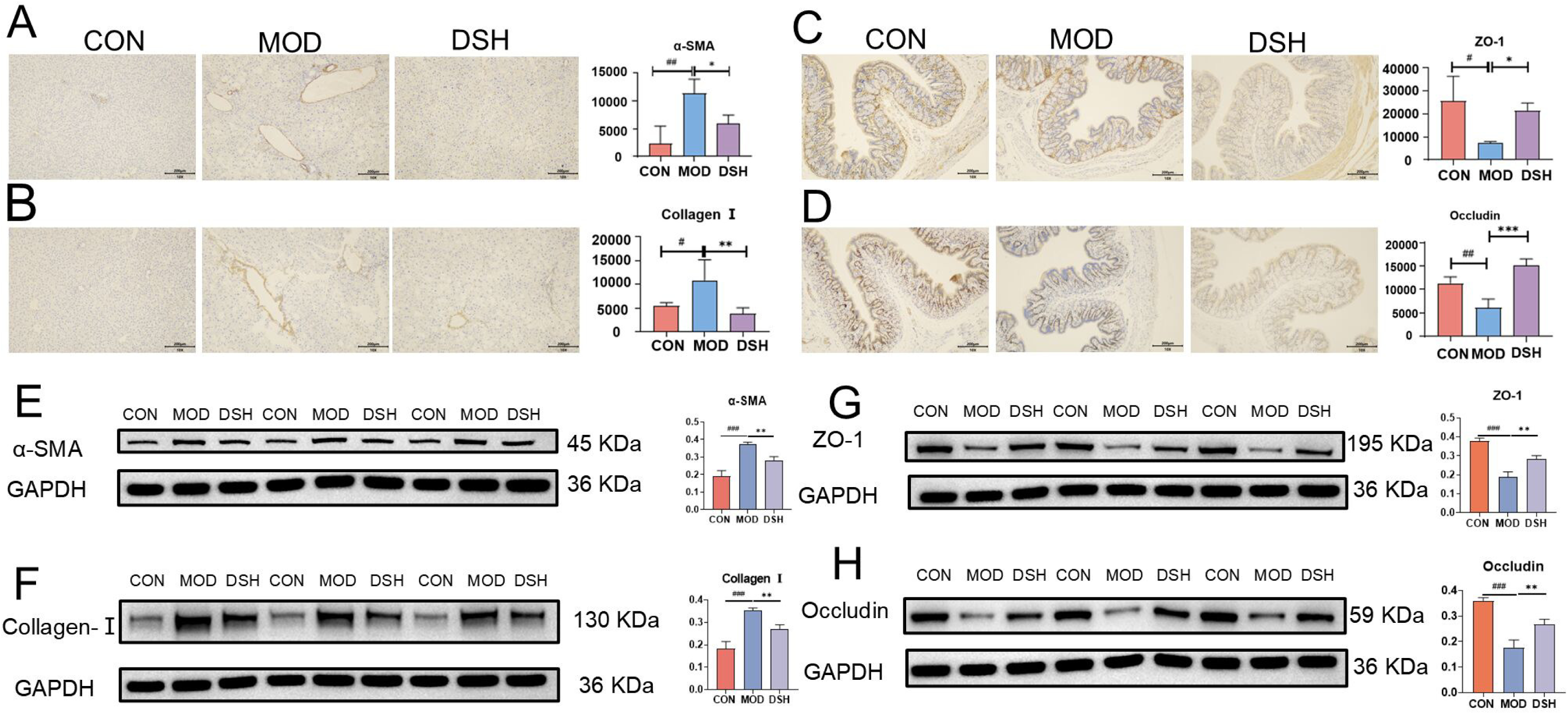
Immunohistochemistry and Immunofluorescence Analysis of Liver and Colon Tissues A:Immunohistochemical staining and quantitative analysis of α-SMA protein in liver tissue. B: Immunohistochemical staining and quantitative analysis of Collagen I protein in liver tissue. C: Immunohistochemical staining and quantitative analysis of ZO-1 protein in colon tissue. D: Immunohistochemical staining and quantitative analysis of Occludin protein in colon tissue; E, F: Detection of α-SMA and Collagen I expression in liver tissues of each group using WB analysis; G, H: Examination of ZO-1 and Occludin expression in colonic tissues of each group by WB method Compare with MOD:\*\*\**p*<0.001,\*\**p*<0.01,\**p*<0.05; Compare with CON:###*p*< 0.001,##*p*<0.01,#*p*<0.05

**Fig. 3.**
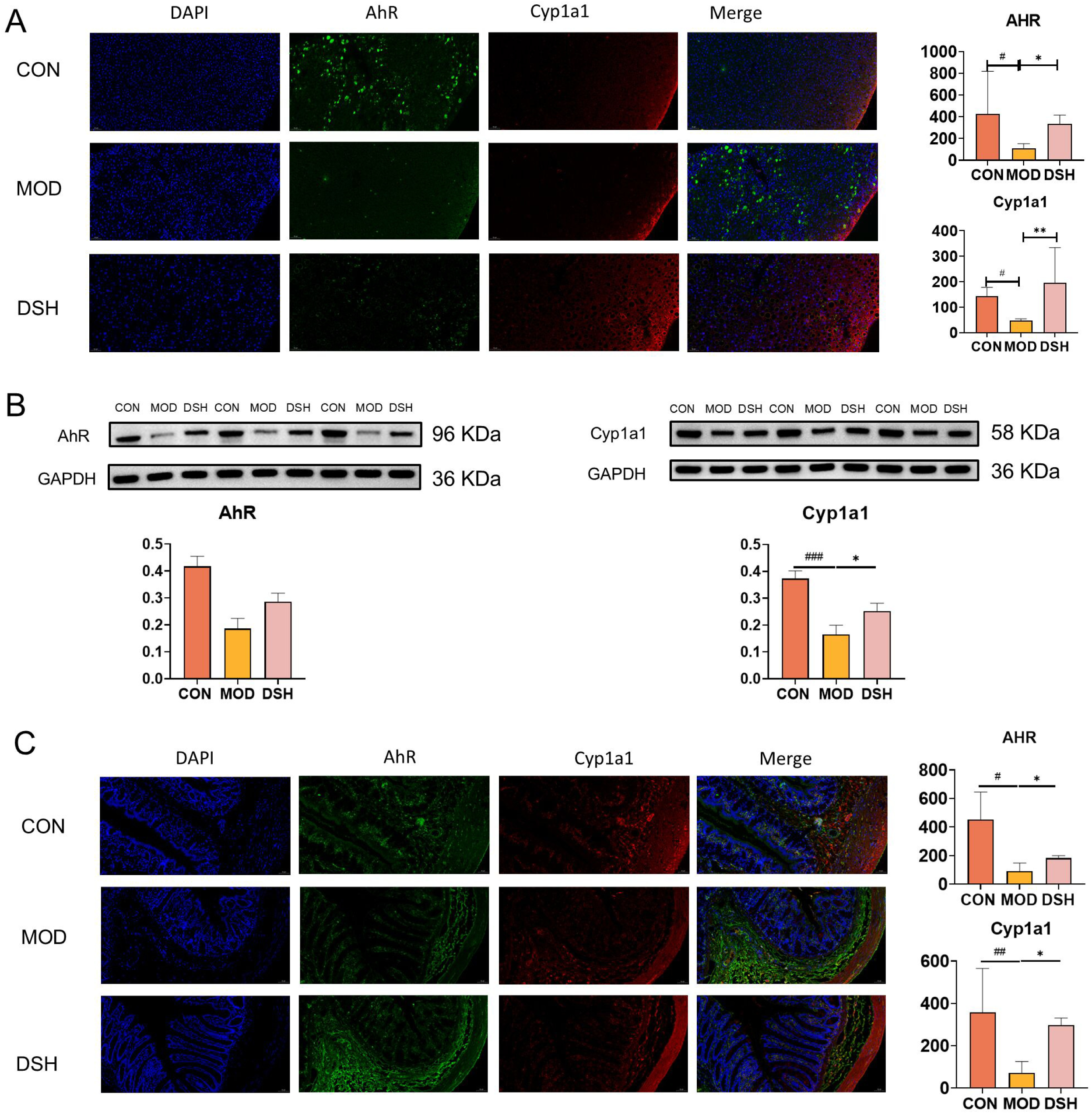
Analysis of AhR and Cyp1a1 Expression in Liver and Colon Tissues A: Immunofluorescence analysis of AhR and Cyp1a1 in liver tissue; B: WB detection of AhR and Cyp1a1 expression in liver tissue; C: Immunofluorescence analysis of AhR and Cyp1a1 in colon tissue. Compare with MOD:\*\*\**p*<0.001,\*\**p*<0.01,\**p*<0.05; Compare with CON:###*p*< 0.001,##*p*<0.01,#*p*<0.05

The impact of the ALI model and DSH intervention on the protein expression of AhR and its downstream target gene Cyp1a1 in liver and colonic tissues was evaluated using immunofluorescence technique. Based on serum biochemical indicators and pathological section results, the DSH group demonstrated the most effective improvement in ALI. Therefore, we conducted immunofluorescence analysis on the liver(Fig 3 A) and colonic(Fig 3 C) tissues of DSH. The results indicated that in the CON group, strong fluorescent signals for AhR and Cyp1a1 were observed in liver tissue, while a clear expression pattern of AhR and Cyp1a1 was also present in colonic tissue. In the MOD group, the fluorescence intensity of AhR and Cyp1a1 in both liver and colonic tissues was significantly reduced compared to the CON group. However, following DSH treatment, the protein expression of AhR and Cyp1a1 in liver and colonic tissues was significantly restored, with the fluorescence intensity of Cyp1a1 returning to near-normal levels. Additionally, the expression of AhR and Cyp1a1 in colonic mucosa was also markedly enhanced. Furthermore, WB analysis demonstrated that ALI reduced the expression of AhR and Cyp1a1 in liver tissue, while DSH treatment led to an increase in the expression levels of AhR and Cyp1a1(Fig 3 B).

### 3.5 Results of Untargeted Serum Metabolomics

#### Analysis of Serum Metabolic Profile

The obtained serum metabolic components were analyzed using EZinfo 3.0 for Waters. Based on pathological and biochemical indicator results, we observed that the DSH group exhibited the best recovery effect on ALI. Therefore, we conducted a metabolomic study on the DSH group. We employed PCA and PLS-DA (Fig 4 A, B) to reduce the dimensionality of the data obtained from the CON, MOD, Sily, DSH, and QC groups. The high clustering of QC samples in the PCA plot indicated the stability and reproducibility of the instrumentation and data acquisition methods used in this experiment. The metabolites confirmed the differential relationships among the various groups. To better interpret the model, we further analyzed the normal group and the model group using OPLS-DA (Figure 4 C, D, E). The samples from different groups could be completely separated on the OPLS-DA plot, indicating good model fit and predictive ability. The R2 and Q2 values in positive mode were 0.9764 and 0.0516, respectively, while in negative mode, they were 0.9957 and 0.3499, respectively.

**Fig 4.**
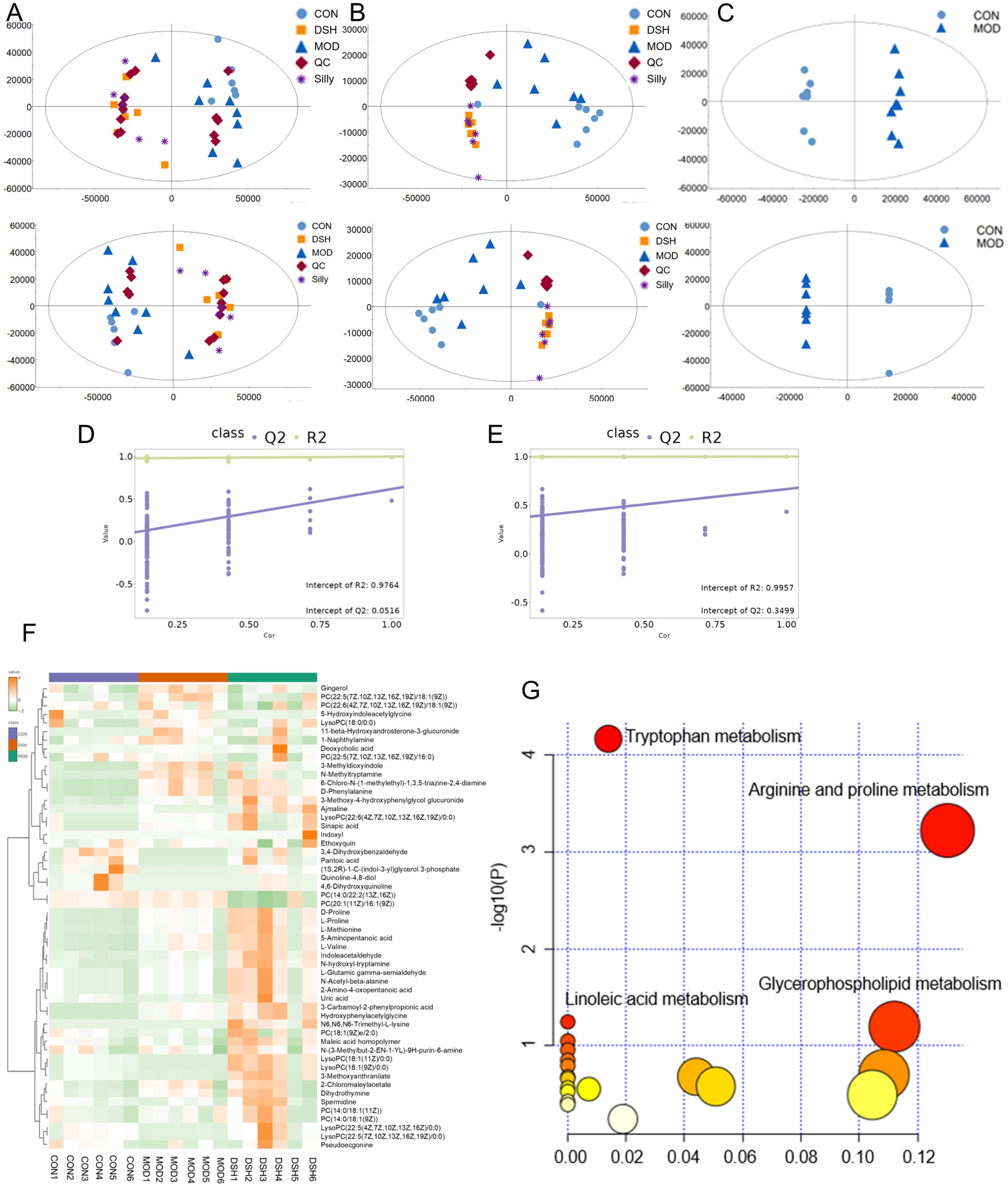
Non-targeted metabolomics study of DS on ALI rat serum. A: PCA analysis in positive and negative ion modes; B: PLS-DA analysis in positive and negative ion modes; C: OPLS-DA analysis in positive and negative ion modes; D, E: Validation analysis in positive and negative ion modes; F: Cluster analysis of non-targeted differential components in serum; G: Pathway analysis of non-targeted differential components in serum.

### 3.6 Screening of Differential Metabolites

To better elucidate the differential relationships within the model, we conducted a further analysis of the differential metabolites among the groups by verifying the disparities in serum components. We employed Mass Frontiers software for the analysis and secondary identification of the screened differential metabolites. To mitigate the risk of false positives or model overfitting that may arise from a single statistical approach, we adopted a comprehensive methodology to determine the differential metabolites. This encompassed a t-test with *p* < 0.05 and VIP > 1 in univariate analysis. Through our analysis, we identified a total of 36 differential metabolites (Table 2).

**Tab 2.**
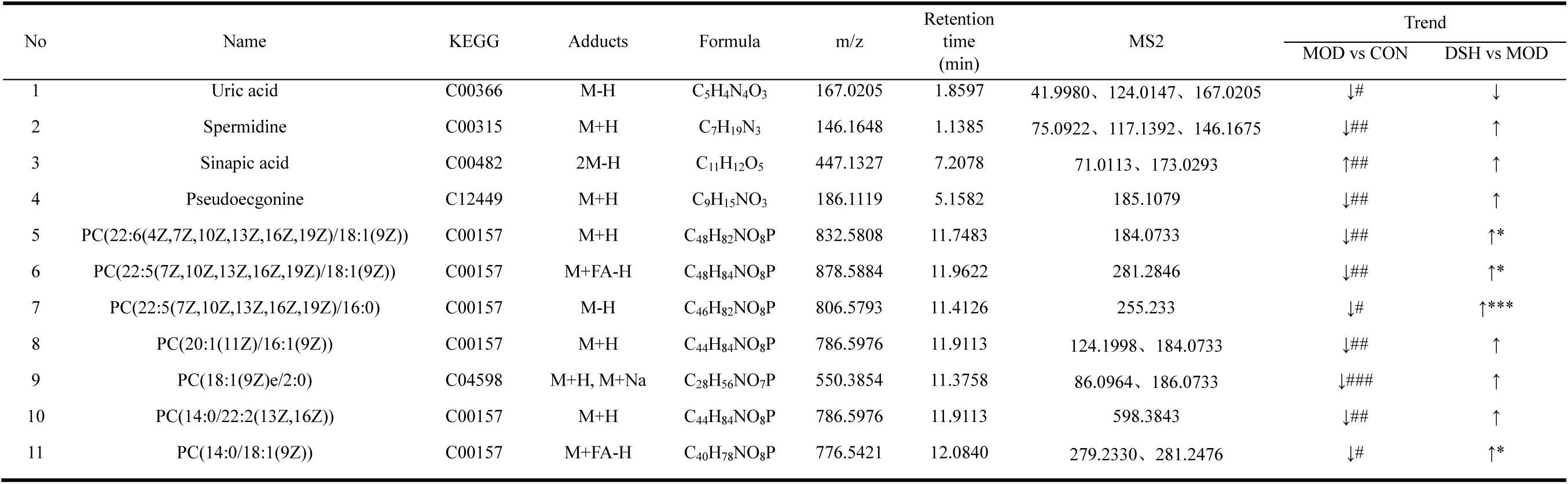
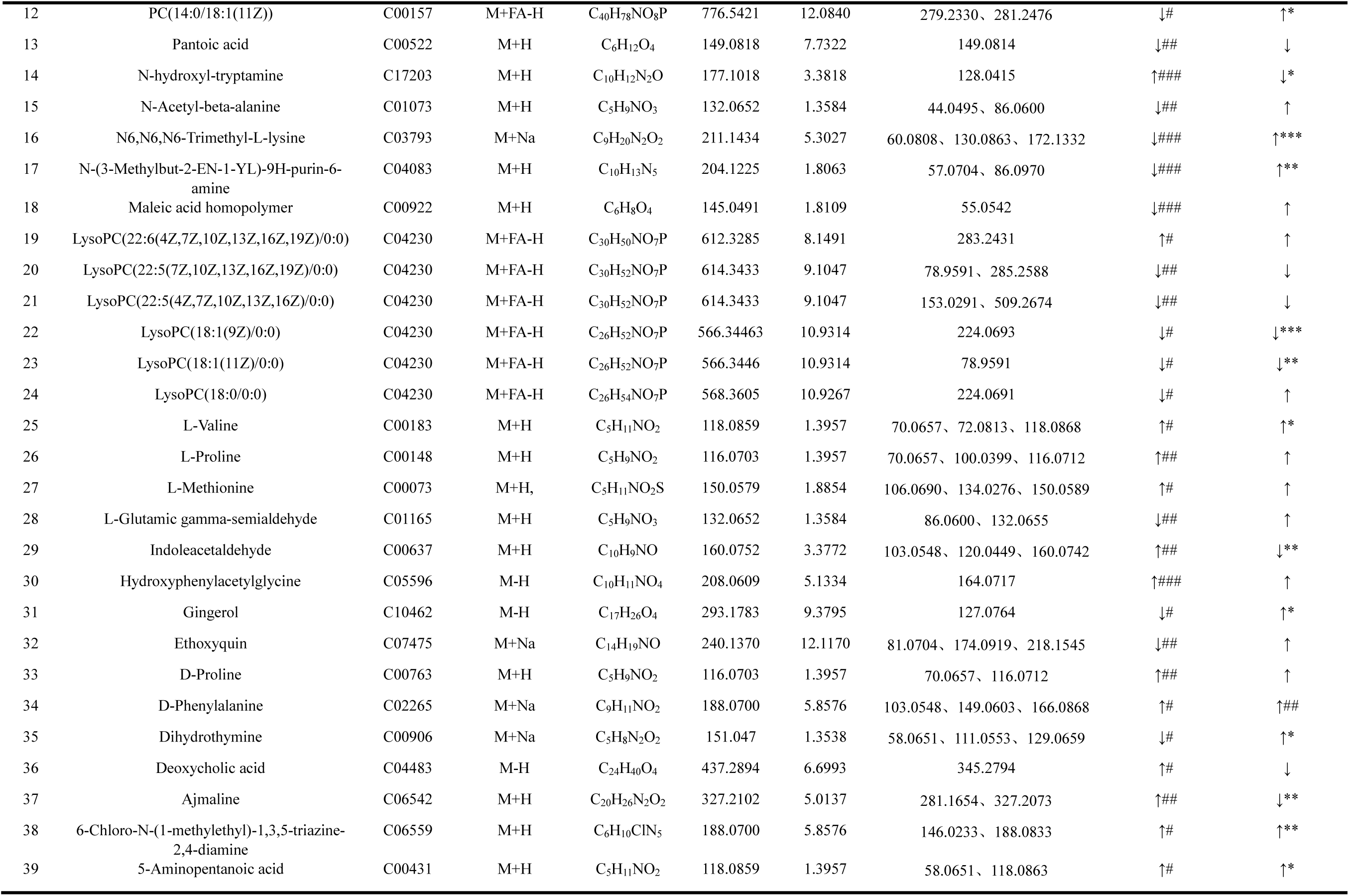
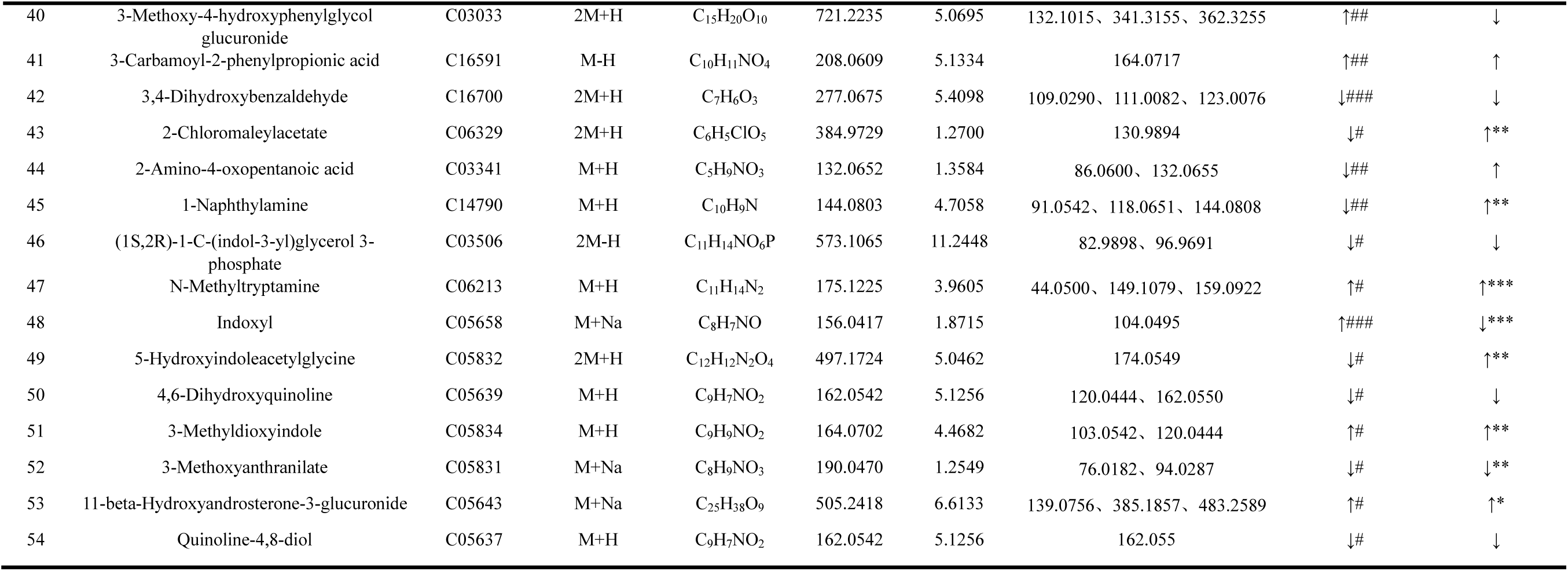
Differential metabolic components in the serum of ALI rats treated with DS.

| No | Name | KEGG | Adducts | Formula | m/z | Retention time (min) | MS2 | Trend |  |
| --- | --- | --- | --- | --- | --- | --- | --- | --- | --- |
|  |  |  |  |  |  |  |  | MOD vs CON | DSH vs MOD |
| 1 | Uric acid | C00366 | M-H | C<sub>5</sub>H<sub>4</sub>N<sub>4</sub>O<sub>3</sub> | 167.0205 | 1.8597 | 41.9980、124.0147、167.0205 | ↓# | ↓ |
| 2 | Spermidine | C00315 | M+H | C<sub>7</sub>H<sub>19</sub>N<sub>3</sub> | 146.1648 | 1.1385 | 75.0922、117.1392、146.1675 | ↓## | ↑ |
| 3 | Sinapic acid | C00482 | 2M-H | C<sub>11</sub>H<sub>12</sub>O<sub>5</sub> | 447.1327 | 7.2078 | 71.0113、173.0293 | ↑## | ↑ |
| 4 | Pseudoecgonine | C12449 | M+H | C<sub>9</sub>H<sub>15</sub>NO<sub>3</sub> | 186.1119 | 5.1582 | 185.1079 | ↓## | ↑ |
| 5 | PC(22:6(4Z,7Z,10Z,13Z,16Z,19Z)/18:1(9Z)) | C00157 | M+H | C<sub>48</sub>H<sub>82</sub>NO<sub>8</sub>P | 832.5808 | 11.7483 | 184.0733 | ↓## | ↑* |
| 6 | PC(22:5(7Z,10Z,13Z,16Z,19Z)/18:1(9Z)) | C00157 | M+FA-H | C<sub>48</sub>H<sub>84</sub>NO<sub>8</sub>P | 878.5884 | 11.9622 | 281.2846 | ↓## | ↑* |
| 7 | PC(22:5(7Z,10Z,13Z,16Z,19Z)/16:0) | C00157 | M-H | C<sub>46</sub>H<sub>82</sub>NO<sub>8</sub>P | 806.5793 | 11.4126 | 255.233 | ↓# | ↑*** |
| 8 | PC(20:1(11Z)/16:1(9Z)) | C00157 | M+H | C<sub>44</sub>H<sub>84</sub>NO<sub>8</sub>P | 786.5976 | 11.9113 | 124.1998、184.0733 | ↓## | ↑ |
| 9 | PC(18:1(9Z)e/2:0) | C04598 | M+H, M+Na | C<sub>28</sub>H<sub>56</sub>NO<sub>7</sub>P | 550.3854 | 11.3758 | 86.0964、186.0733 | ↓### | ↑ |
| 10 | PC(14:0/22:2(13Z,16Z)) | C00157 | M+H | C<sub>44</sub>H<sub>84</sub>NO<sub>8</sub>P | 786.5976 | 11.9113 | 598.3843 | ↓## | ↑ |
| 11 | PC(14:0/18:1(9Z)) | C00157 | M+FA-H | C<sub>40</sub>H<sub>78</sub>NO<sub>8</sub>P | 776.5421 | 12.0840 | 279.2330、281.2476 | ↓# | ↑* |
| 12 | PC(14:0/18:1(11Z)) | C00157 | M+FA-H | C<sub>40</sub>H<sub>78</sub>NO<sub>8</sub>P | 776.5421 | 12.0840 | 279.2330、281.2476 | ↓# | ↑* |
| 13 | Pantoic acid | C00522 | M+H | C<sub>6</sub>H<sub>12</sub>O<sub>4</sub> | 149.0818 | 7.7322 | 149.0814 | ↓## | ↓ |
| 14 | N-hydroxyl-tryptamine | C17203 | M+H | C<sub>10</sub>H<sub>12</sub>N<sub>2</sub>O | 177.1018 | 3.3818 | 128.0415 | ↑### | ↓* |
| 15 | N-Acetyl-beta-alanine | C01073 | M+H | C<sub>5</sub>H<sub>9</sub>NO<sub>3</sub> | 132.0652 | 1.3584 | 44.0495、86.0600 | ↓## | ↑ |
| 16 | N6,N6,N6-Trimethyl-L-lysine | C03793 | M+Na | C<sub>9</sub>H<sub>20</sub>N<sub>2</sub>O<sub>2</sub> | 211.1434 | 5.3027 | 60.0808、130.0863、172.1332 | ↓### | ↑*** |
| 17 | N-(3-Methylbut-2-EN-1-YL)-9H-purin-6-amine | C04083 | M+H | C<sub>10</sub>H<sub>13</sub>N<sub>5</sub> | 204.1225 | 1.8063 | 57.0704、86.0970 | ↓### | ↑** |
| 18 | Maleic acid homopolymer | C00922 | M+H | C<sub>6</sub>H<sub>8</sub>O<sub>4</sub> | 145.0491 | 1.8109 | 55.0542 | ↓### | ↑ |
| 19 | LysoPC(22:6(4Z,7Z,10Z,13Z,16Z,19Z)/0:0) | C04230 | M+FA-H | C<sub>30</sub>H<sub>50</sub>NO<sub>7</sub>P | 612.3285 | 8.1491 | 283.2431 | ↑# | ↑ |
| 20 | LysoPC(22:5(7Z,10Z,13Z,16Z,19Z)/0:0) | C04230 | M+FA-H | C<sub>30</sub>H<sub>52</sub>NO<sub>7</sub>P | 614.3433 | 9.1047 | 78.9591、285.2588 | ↓## | ↓ |
| 21 | LysoPC(22:5(4Z,7Z,10Z,13Z,16Z)/0:0) | C04230 | M+FA-H | C<sub>30</sub>H<sub>52</sub>NO<sub>7</sub>P | 614.3433 | 9.1047 | 153.0291、509.2674 | ↓## | ↓ |
| 22 | LysoPC(18:1(9Z)/0:0) | C04230 | M+FA-H | C<sub>26</sub>H<sub>52</sub>NO<sub>7</sub>P | 566.34463 | 10.9314 | 224.0693 | ↓# | ↓*** |
| 23 | LysoPC(18:1(11Z)/0:0) | C04230 | M+FA-H | C<sub>26</sub>H<sub>52</sub>NO<sub>7</sub>P | 566.3446 | 10.9314 | 78.9591 | ↓# | ↓** |
| 24 | LysoPC(18:0/0:0) | C04230 | M+FA-H | C<sub>26</sub>H<sub>54</sub>NO<sub>7</sub>P | 568.3605 | 10.9267 | 224.0691 | ↓# | ↑ |
| 25 | L-Valine | C00183 | M+H | C<sub>5</sub>H<sub>11</sub>NO<sub>2</sub> | 118.0859 | 1.3957 | 70.0657、72.0813、118.0868 | ↑# | ↑* |
| 26 | L-Proline | C00148 | M+H | C<sub>5</sub>H<sub>9</sub>NO<sub>2</sub> | 116.0703 | 1.3957 | 70.0657、100.0399、116.0712 | ↑## | ↑ |
| 27 | L-Methionine | C00073 | M+H, | C<sub>5</sub>H<sub>11</sub>NO<sub>2</sub>S | 150.0579 | 1.8854 | 106.0690、134.0276、150.0589 | ↑# | ↑ |
| 28 | L-Glutamic gamma-semialdehyde | C01165 | M+H | C<sub>5</sub>H<sub>9</sub>NO<sub>3</sub> | 132.0652 | 1.3584 | 86.0600、132.0655 | ↓## | ↑ |
| 29 | Indoleacetaldehyde | C00637 | M+H | C<sub>10</sub>H<sub>9</sub>NO | 160.0752 | 3.3772 | 103.0548、120.0449、160.0742 | ↑## | ↓** |
| 30 | Hydroxyphenylacetyl glycine | C05596 | M-H | C<sub>10</sub>H<sub>11</sub>NO<sub>4</sub> | 208.0609 | 5.1334 | 164.0717 | ↑### | ↑ |
| 31 | Gingerol | C10462 | M-H | C<sub>17</sub>H<sub>26</sub>O<sub>4</sub> | 293.1783 | 9.3795 | 127.0764 | ↓# | ↑* |
| 32 | Ethoxyquin | C07475 | M+Na | C<sub>14</sub>H<sub>19</sub>NO | 240.1370 | 12.1170 | 81.0704、174.0919、218.1545 | ↓## | ↑ |
| 33 | D-Proline | C00763 | M+H | C<sub>5</sub>H<sub>9</sub>NO<sub>2</sub> | 116.0703 | 1.3957 | 70.0657、116.0712 | ↑## | ↑ |
| 34 | D-Phenylalanine | C02265 | M+Na | C<sub>9</sub>H<sub>11</sub>NO<sub>2</sub> | 188.0700 | 5.8576 | 103.0548、149.0603、166.0868 | ↑# | ↑## |
| 35 | Dihydrothymine | C00906 | M+Na | C<sub>5</sub>H<sub>8</sub>N<sub>2</sub>O<sub>2</sub> | 151.047 | 1.3538 | 58.0651、111.0553、129.0659 | ↓# | ↑* |
| 36 | Deoxycholic acid | C04483 | M-H | C<sub>24</sub>H<sub>40</sub>O<sub>4</sub> | 437.2894 | 6.6993 | 345.2794 | ↑# | ↓ |
| 37 | Ajmaline | C06542 | M+H | C<sub>20</sub>H<sub>26</sub>N<sub>2</sub>O<sub>2</sub> | 327.2102 | 5.0137 | 281.1654、327.2073 | ↑## | ↓** |
| 38 | 6-Chloro-N-(1-methylethyl)-1,3,5-triazine-2,4-diamine | C06559 | M+H | C<sub>6</sub>H<sub>10</sub>ClN<sub>5</sub> | 188.0700 | 5.8576 | 146.0233、188.0833 | ↑# | ↑** |
| 39 | 5-Aminopentanoic acid | C00431 | M+H | C<sub>5</sub>H<sub>11</sub>NO<sub>2</sub> | 118.0859 | 1.3957 | 58.0651、118.0863 | ↑# | ↑* |
| 40 | 3-Methoxy-4-hydroxyphenylglycol glucuronide | C03033 | 2M+H | C<sub>15</sub>H<sub>20</sub>O<sub>10</sub> | 721.2235 | 5.0695 | 132.1015、341.3155、362.3255 | ↑## | ↓ |
| 41 | 3-Carbamoyl-2-phenylpropionic acid | C16591 | M-H | C<sub>10</sub>H<sub>11</sub>NO<sub>4</sub> | 208.0609 | 5.1334 | 164.0717 | ↑## | ↑ |
| 42 | 3,4-Dihydroxybenzaldehyde | C16700 | 2M+H | C<sub>7</sub>H<sub>6</sub>O<sub>3</sub> | 277.0675 | 5.4098 | 109.0290、111.0082、123.0076 | ↓### | ↓ |
| 43 | 2-Chloromaleylacetate | C06329 | 2M+H | C<sub>6</sub>H<sub>5</sub>ClO<sub>5</sub> | 384.9729 | 1.2700 | 130.9894 | ↓# | ↑** |
| 44 | 2-Amino-4-oxopentanoic acid | C03341 | M+H | C<sub>5</sub>H<sub>9</sub>NO<sub>3</sub> | 132.0652 | 1.3584 | 86.0600、132.0655 | ↓## | ↑ |
| 45 | 1-Naphthylamine | C14790 | M+H | C<sub>10</sub>H<sub>9</sub>N | 144.0803 | 4.7058 | 91.0542、118.0651、144.0808 | ↓## | ↑** |
| 46 | (1S,2R)-1-C-(indol-3-yl)glycerol 3-phosphate | C03506 | 2M-H | C<sub>11</sub>H<sub>14</sub>NO<sub>6</sub>P | 573.1065 | 11.2448 | 82.9898、96.9691 | ↓# | ↓ |
| 47 | N-Methyltryptamine | C06213 | M+H | C<sub>11</sub>H<sub>14</sub>N<sub>2</sub> | 175.1225 | 3.9605 | 44.0500、149.1079、159.0922 | ↑# | ↑*** |
| 48 | Indoxyl | C05658 | M+Na | C<sub>8</sub>H<sub>7</sub>NO | 156.0417 | 1.8715 | 104.0495 | ↑### | ↓*** |
| 49 | 5-Hydroxyindoleacetyl glycine | C05832 | 2M+H | C<sub>12</sub>H<sub>12</sub>N<sub>2</sub>O<sub>4</sub> | 497.1724 | 5.0462 | 174.0549 | ↓# | ↑** |
| 50 | 4,6-Dihydroxyquinoline | C05639 | M+H | C<sub>9</sub>H<sub>7</sub>NO<sub>2</sub> | 162.0542 | 5.1256 | 120.0444、162.0550 | ↓# | ↓ |
| 51 | 3-Methyldioxyindole | C05834 | M+H | C<sub>9</sub>H<sub>9</sub>NO<sub>2</sub> | 164.0702 | 4.4682 | 103.0542、120.0444 | ↑# | ↑** |
| 52 | 3-Methoxyanthranilate | C05831 | M+Na | C<sub>8</sub>H<sub>9</sub>NO<sub>3</sub> | 190.0470 | 1.2549 | 76.0182、94.0287 | ↓# | ↓** |
| 53 | 11-beta-Hydroxyandrosterone-3-glucuronide | C05643 | M+Na | C<sub>25</sub>H<sub>38</sub>O<sub>9</sub> | 505.2418 | 6.6133 | 139.0756、385.1857、483.2589 | ↑# | ↑* |
| 54 | Quinoline-4,8-diol | C05637 | M+H | C<sub>9</sub>H<sub>7</sub>NO<sub>2</sub> | 162.0542 | 5.1256 | 162.055 | ↓# | ↓ |

### 3.7 Clustering Analysis of Differential Metabolites in Serum

The heatmap was generated using the Wekemo Bioincloud platform.The clustering heatmap can illustrate the clustering patterns of different metabolites among samples. Clustering heatmap analysis of the selected differential metabolites was performed using MetaboAnalyst 6.0 (Fig 4 F). The x-axis represents the groups, while the y-axis represents the differential metabolites. From the clustering heatmap, it can be observed that MOD is distant from all other groups, CON exhibits good clustering, and DSH is close to CON. This suggests that DS may have a positive effect in the treatment of ALI.

### 3.8 Impact of DS on Metabolic Pathways in ALI Rats

Enrichment pathway analysis enables the examination of differential metabolite components (Fig 4 G). Utilizing the MetaboAnalyst 6.0 platform, potential target pathways were identified and visually represented through data visualization. Among these, several profound metabolic pathways emerged, including Tryptophan metabolism、 Arginine and proline metabolism、Glycerophospholipid metabolism、Linoleic acid metabolism.

### 3.9 Targeted Metabolomics Analysis of Tryptophan

Based on our analysis of non-targeted differential metabolite pathways and related studies, we hypothesized that ALI might be associated with the tryptophan metabolism pathway. Therefore, we conducted a tryptophan-targeted metabolomics analysis on fecal samples. In this analysis, we used 51 standard substances, including the final products of tryptophan metabolism such as xanthurenic acid, shikimic acid, L-tryptophan, and indole, as controls(S1). The results revealed eight tryptophan-related differential metabolites between the CON and MOD groups (Fig 5 A) (*p* < 0.05). Following DSH treatment, some tryptophan metabolites returned to normal levels. Among them, 3-(3-Indolyl)-2-oxopropanoic acid, 4-Hydroxyphenylacetic acid, and L-Kynurenine were identified as common differential metabolites across all experimental groups. Based on the results of the targeted metabolomics analysis, we speculated that ALI might be regulated by the tryptophan metabolism pathway.

**Fig 5.**
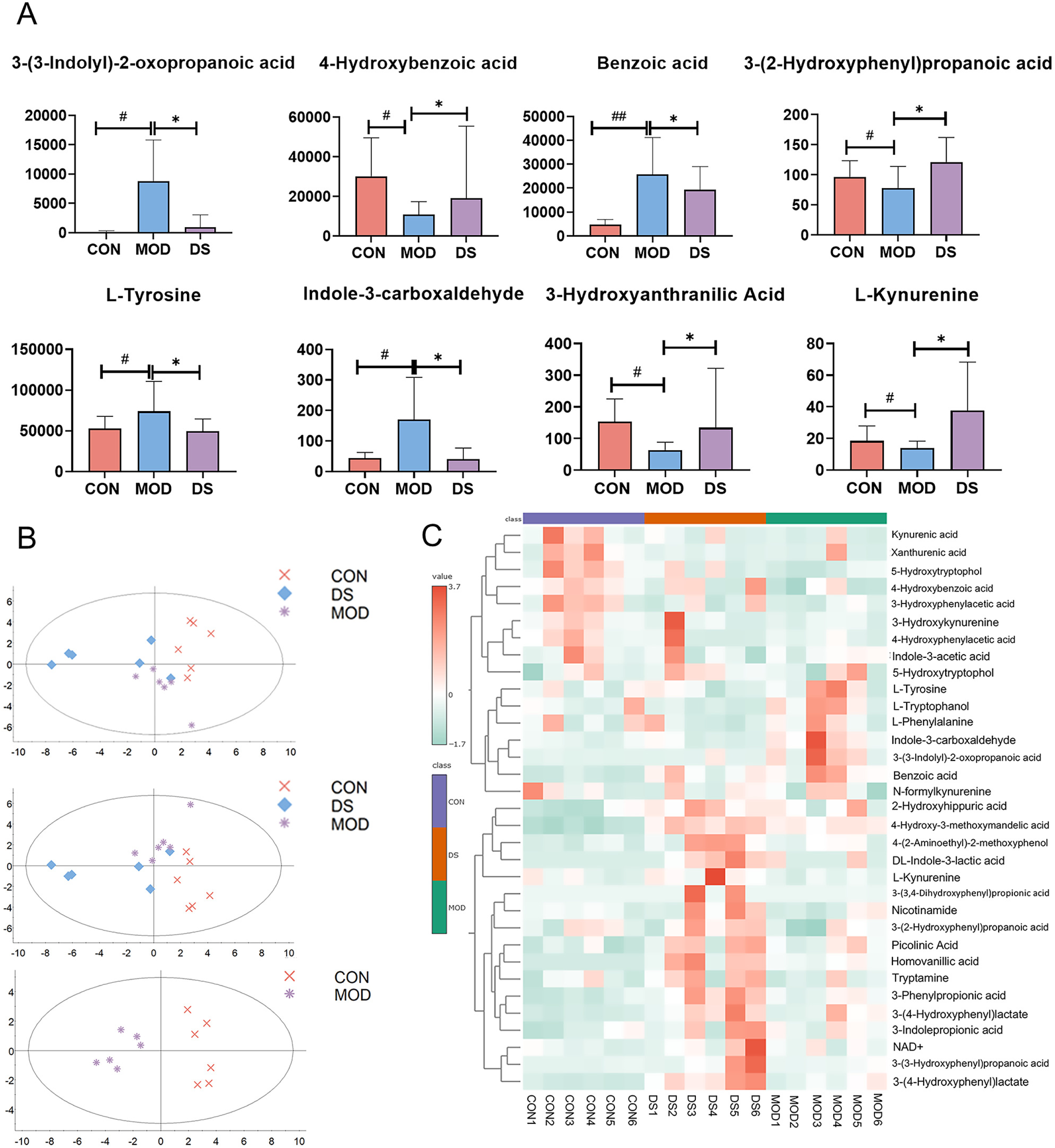
Tryptophan Targeted Metabolomics Metabolic Profiling and Differential Metabolic Component Cluster Analysis (A) Differential metabolites of tryptophan,(B) PCA, PLS-DA, and OPLS-DA analysis of tryptophan targeted metabolism,(C) Cluster analysis of differential metabolites in tryptophan targeted metabolism. Compare with MOD:\*\*\**p*<0.001,\*\**p*<0.01,\**p*<0.05; Compare with CON:###*p*< 0.001,##*p*<0.01,#*p*<0.05

### 3.10 Metabolic Profiling Analysis and Metabolic Heatmap

According to the PCA, PLS-DA, and OPLS-DA plots of tryptophan metabolites (Figure 5 B), there was a significant separation between MOD and CON, while CON and DSH showed substantial overlap. The OPLS-DA results indicated a clear separation between CON and MOD, suggesting that DSH could restore differential metabolites by influencing the tryptophan metabolism pathway, thereby exerting a therapeutic effect on ALI. The targeted metabolomics heatmap (Fig 5 C) further supported this inference, showing a closer relationship between CON and DSH, indicating DS potential therapeutic role in ALI

### 3.11 DS ameliorates ALI by modulating gut microbiota

The gut microbiota composition of four mouse groups was analyzed using 16S rRNA gene sequencing technology. Alpha diversity, which provides insights into the complexity of species diversity within samples, was calculated using QIIME2 software. Four metrics were evaluated in this study, including Chao1, observed_otus, Shannon, and Simpson, which collectively capture the intricacies of species diversity, enabling a more accurate and comprehensive assessment of sample diversity. Beta diversity was also computed using QIIME2, and visualizations were created using the R package (v3.5.2). Sequence alignment was performed using Blast, and representative sequences were annotated based on the SILVA database.

Alpha diversity serves as a measure of microbial richness and evenness within samples. As shown in Fig 6 A, compared to MOD, the DSH and Sily groups demonstrated significant increases in these diversity indices. Conversely, DS and Sliy groups exhibited significant reductions in observed_otus, Chao1, Shannon, and Simpson indices when compared to MOD. These findings suggest that DSH and Sily have similar effects, significantly reducing the diversity and richness of the gut microbiota in ALI rats

**Fig 6.**
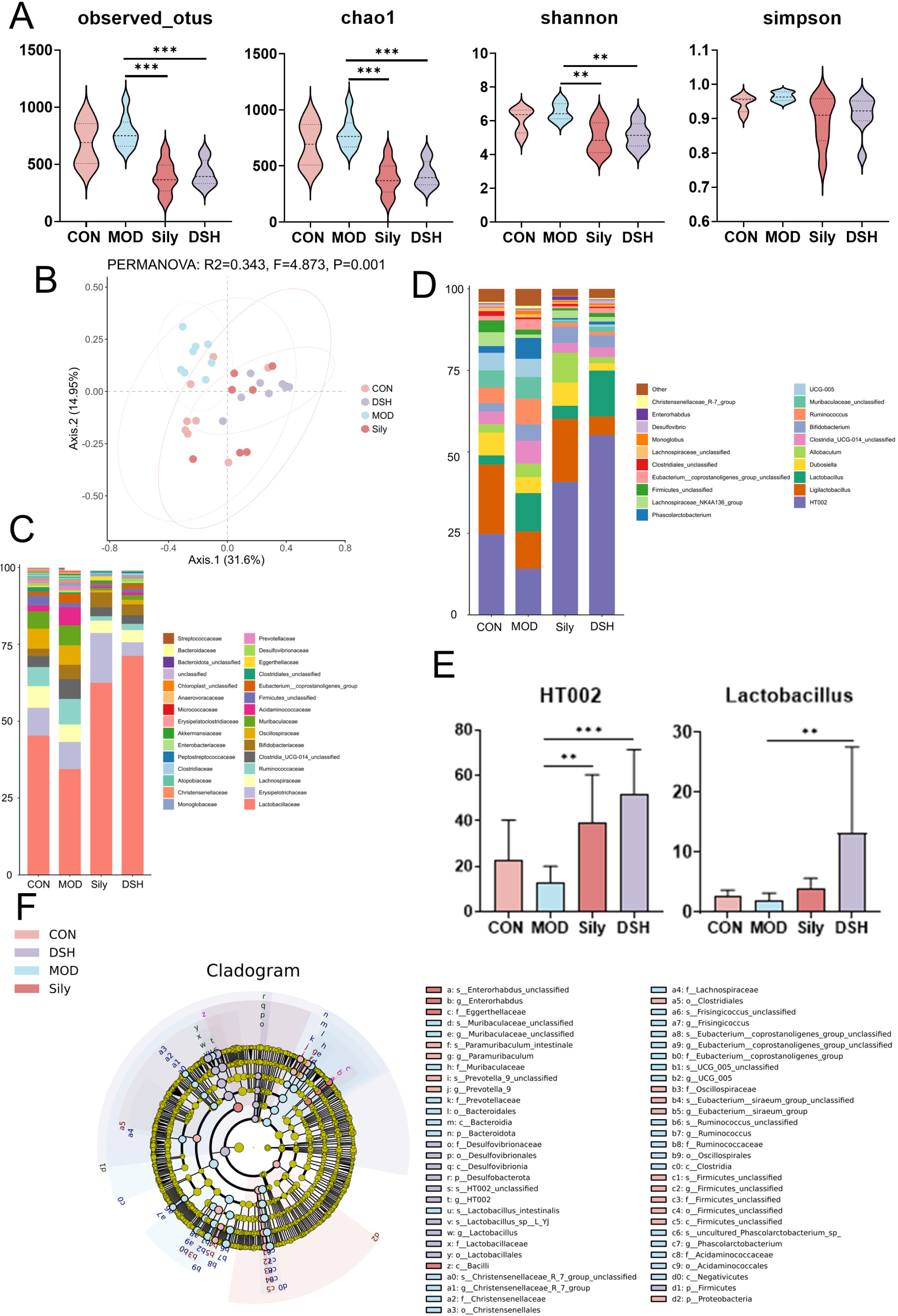
Analysis of the effect of DS on the gut microbiota of ALI rats. A: Alpha diversity analysis; B: Beta diversity analysis; C: Changes in gut microbiota at the phylum level; D, E: Changes in gut microbiota at the genus level; F: LEfSe analysis. Compare with MOD:\*\*\**p*<0.001,\*\**p*<0.01,\**p*<0.05; Compare with CON:###*p*< 0.001,##*p*<0.01,#*p*<0.05

In the study of biological communities, beta diversity is often used to evaluate the degree of difference between species communities by measuring the distance between them. The beta diversity among various groups is illustrated in (Fig 6 B). The MOD group is relatively distant from the other treatment groups, while the DS group and the Sliy group are closer to each other. This suggests that after administration of the medication, the effects of DS and Sliy are similar, resulting in smaller differences in their microbial community composition.

Further evaluation of the species composition was performed for each group. At the phylum level, as shown in Fig 6 C, the prominent bacterial families across all groups were *Lactobacillaceae*, *Erysipelotrichaceae*, and *Lachnospiraceae*. Specifically, in ALI rats, the abundance of *Lactobacillaceae* was reduced compared to CON rats, while it was increased after administration of both Sily and DSH compared to ALI rats. At the genus level, as illustrated in Fig 6 D, the relevant microorganisms in each treatment group were primarily represented by *HT002*, *Ligilactobacillus*, and *Lactobacillus*, all of which returned to normal levels after treatment with DS and Sily. Notably, DS and Sily exhibited similar effects on *HT002* and *Lactobacillus* (Fig 6 E), suggesting that their similarity may be reflected in their comparable impacts on these two bacterial genera, both demonstrating improvement in ALI.

Fig 6 F illustrates the significantly abundant species identified by LEfSe analysis at the levels of phylum, class, order, family, and genus. With an LDA score threshold of 3.5, a greater number of species, totaling 30, were enriched in the MOD microbiota, including *c Clostridia*, *o Oscillospirales*, and *p Bacteroidota*. Following the administration of Sily in ALI, four dominant microbiota species underwent enrichment, namely *c_Bacilli*, *g Enterorhabdus*, and *f Eggerthellaceae*. However, in ALI rats treated with DSH, a total of 11 dominant microbiota species were enriched, such as *g HT002*, *g Lactobacillus*, and *p Firmicutes*.

### 3.12 Integrated Analysis of Gut Microbiota and Differential Metabolites

To comprehensively evaluate the therapeutic effects of DS on ALI, we employed Spearman’s correlation analysis to examine the associations between key differential metabolites and significant gut microbiota(Fig 7). Our findings revealed a positive correlation between Lactobacillus and several metabolites, including N-hydroxyl-tryptamine, Indoleacetaldehyde, N-Methyltryptamine, L-Methionine, D-Phenylalanine, 5-Aminopentanoic acid, L-Valine, L-Proline, and D-Proline. Conversely, Lactobacillus was negatively correlated with 3-Methyldioxyindole, Pantoic acid, 4,6-Dihydroxyquinoline, Quinoline-4,8-diol, PC(22:6(4Z,7Z,10Z,13Z,16Z,19Z)/18:1(9Z)), PC(22:5(7Z,10Z,13Z,16Z,19Z)/18:1(9Z)), LysoPC(22:5(7Z,10Z,13Z,16Z,19Z)/0:0), and LysoPC(22:5(4Z,7Z,10Z,13Z,16Z)/0:0

**Fig 7.**
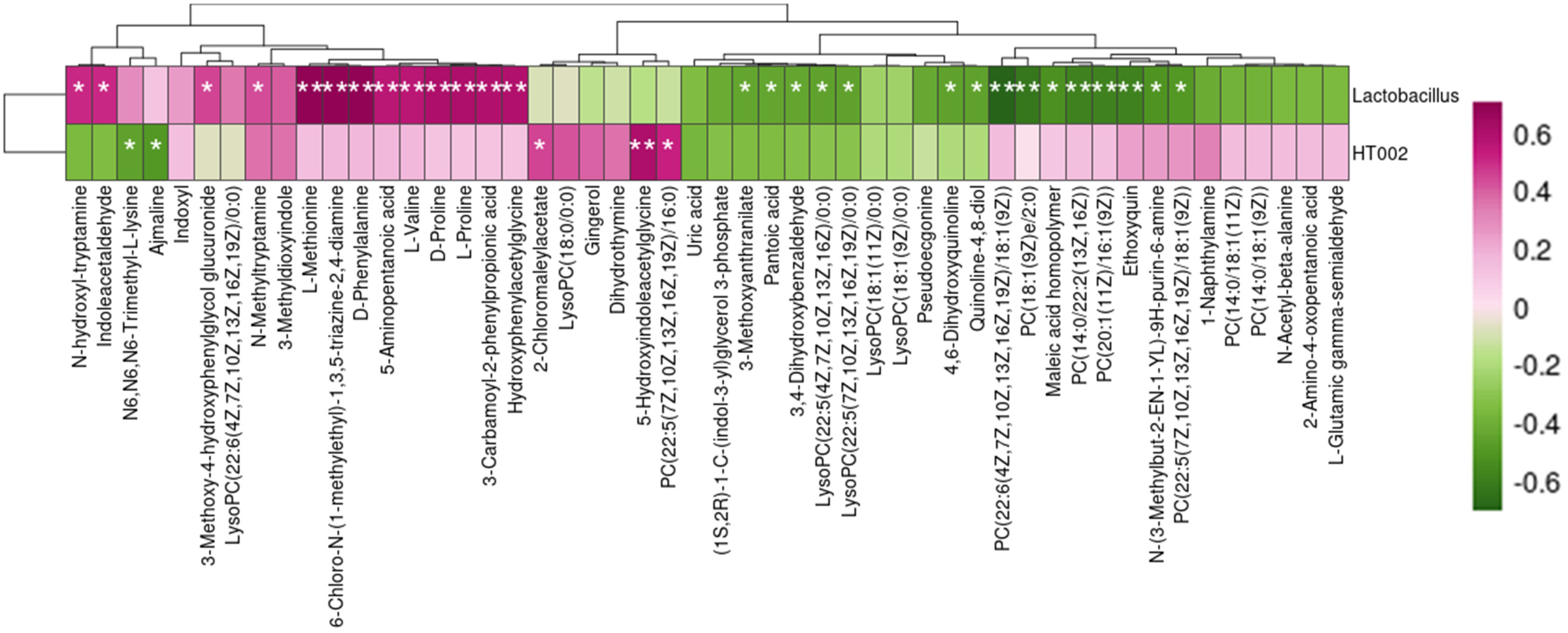
Joint Analysis of Gut Microbiota at the Species Level and Differential Metabolic Components

### 3.13 Therapeutic Effects of FMT on Rats with ALI

#### 3.13.1 DS reduces ALI lesions in pseudo-sterile rats

The staining results of HE (Fig 7 A) and Masson’s trichrome (Fig 7 B) showed significant structural disorder of the liver lobules in ADS, with disordered hepatocyte arrangement, increased vacuolar degeneration and necrosis of cells, and obvious inflammatory cell infiltration within the fibrous septa. Fibrosis was also evident. Quantitative analysis of Masson’s staining further indicated fibrotic lesions with collagen fiber hyperplasia in the ADS group, similar to the characteristics observed in liver tissue sections of AMOD rats, while the liver tissue of AMF rats showed improvement. HE staining of the colon (Fig 7 C) revealed a tightly structured colonic mucosal crypt with abundant goblet cells and no significant inflammation in the lamina propria in the CON group. In AMOD rats, the colonic mucosa showed atrophy, destruction of crypt structure, reduced goblet cell density, and accompanied by massive neutrophil infiltration. Administration of DS fecal microbiota transplantation resulted in mucosal thickness and goblet cell density approaching normal levels, with inflammatory infiltration almost disappearing. The pathological features were highly consistent with those of the CON group. However, no significant improvement was observed in the ADS group.

#### 3.13.2 DS ameliorates ALI in pseudo-germ-free rats by modulating liver fibrosis-related proteins and intestinal barrier function proteins

Imunohistochemical analysis was performed to evaluate α-SMA expression levels. (Fig 8 D) and Collagen-I (Fig 8 E) in liver tissue. Quantitative analysis revealed a significant increase in α-SMA content in the AMOD and ADS groups compared to the CON group, while the α-SMA values in the AMF group tended to be normal. Regarding Collagen-I, as shown in Figure 2 D, there was a notable elevation in its content in the liver tissue of AMOD rats compared to the CON group. However, the Collagen-I content in the liver tissue of AMF rats approached that of the normal group.

**Fig 8.**
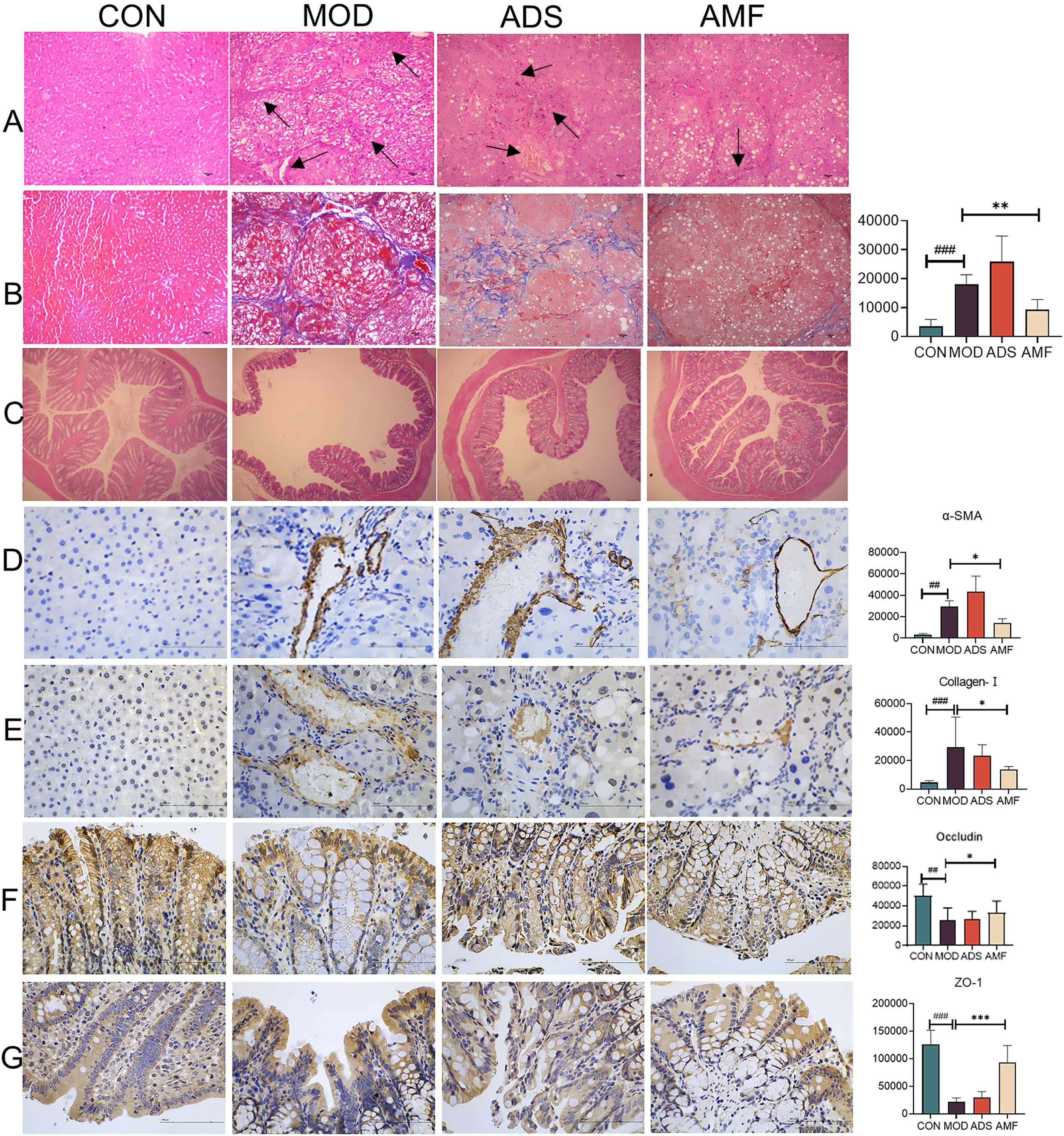
Histopathological study and immunohistochemistry of liver tissue and colon tissue of fecal microbiota transplantation rats. A HE staining of liver tissue (100×); B Masson staining of liver tissue (100×); C: HE staining of colon tissue 100×;D: Immunohistochemistry and quantitative analysis of α-SMA in liver tissue; E: Immunohistochemistry and quantitative analysis of Collagen-Ⅰ in liver tissue; F: Immunohistochemistry and quantitative analysis of Occludin in colon tissue; G: Immunohistochemistry and quantitative analysis of ZO-1 in colon tissue. Compare with AMOD:\*\*\**p*<0.001,\*\**p*<0.01,\**p*<0.05; Compare with CON: ###*p*< 0.001,##*p*<0.01,#*p*<0.05

Furthermore, we performed immunohistochemical staining to visualize the expression of ZO-1 (Fig 8 F) and Occludin (Fig 8 G) in colonic tissue. Compared to the CON group, there was a marked reduction in the positive gene expression of ZO-1 and Occludin in the liver tissue of AMOD rats. Nevertheless, the levels of both ZO-1 and Occludin in the liver tissue of ADS and AMF rats were similar to those of the normal group.

#### 3.13.3 DS fecal bacterial suspension ameliorates ALI in pseudo-germ-free rats by modulating gut microbiota

According to the results of Lefse analysis(Fig 9 A), when setting the LDA value at 3.5, we identified a total of 58 enriched dominant bacterial groups in AMF, including *Lactobacillus*, *Lachnospiraceae*, *Lachnospiraceae*, *Desulfovibrionia*, and others.

**Fig 9.**
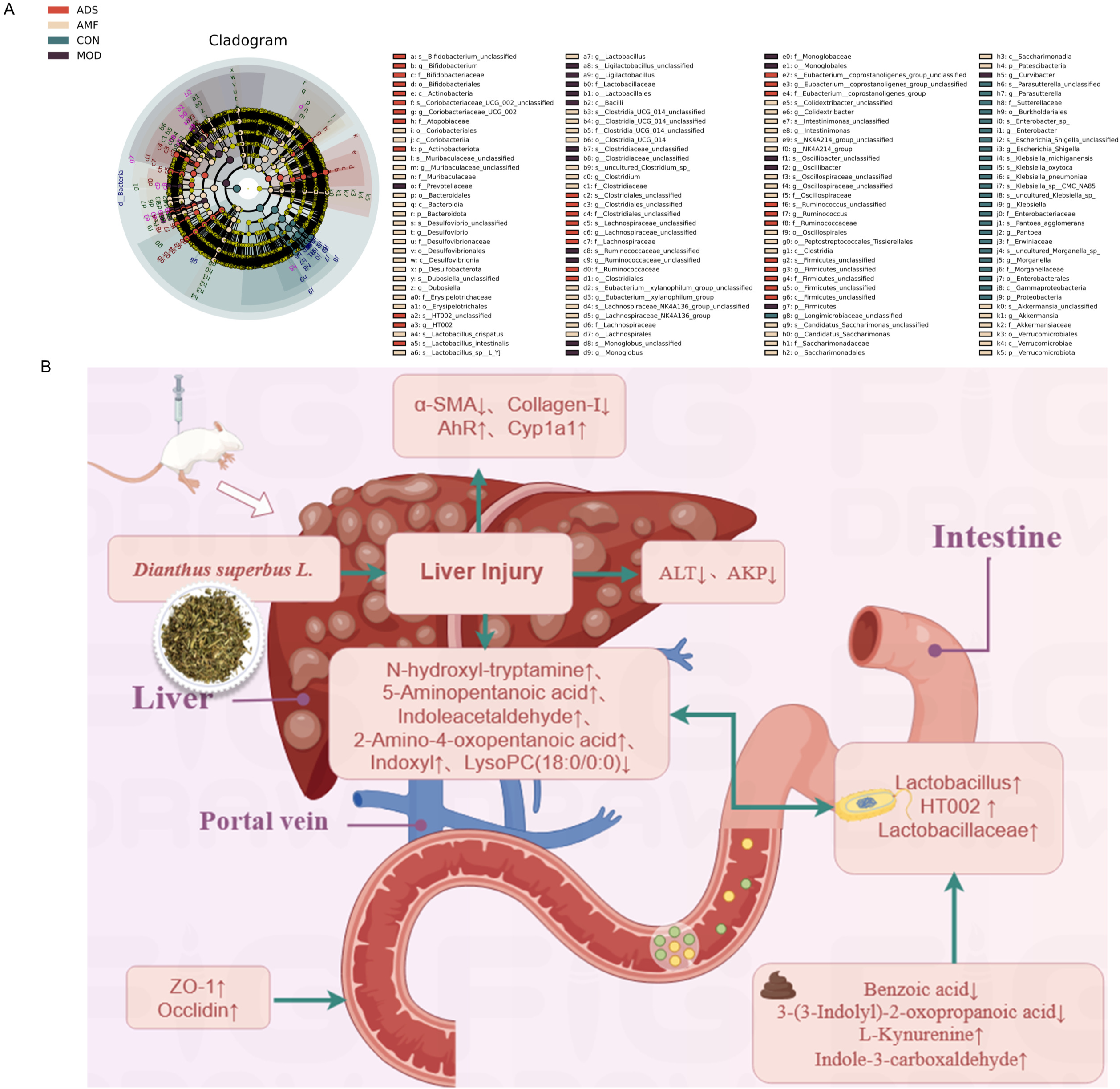
Analysis of gut microbiota in pseudo-germ-free rats and the mechanism of DS action in ALI rats A: Analysis of gut microbiota in pseudo-germ-free rats; B: Mechanism of DS action in ALI rats ↑ indicates upward adjustment, ↓ indicates downward adjustment.

## 4 Discussion

ALI is a common health concern that poses a serious threat to human health due to its wide range of pathogenic factors^31^. There is a growing body of clinical and experimental evidence indicating that the gut microbiota plays a significant role in the pathophysiology of ALI^32^. Manipulation of the gut microbiota is considered a promising strategy for the treatment of ALI. While DS is one of the traditional Mongolian medicines used to treat liver diseases, the mechanism by which it improves ALI remains incompletely understood. Therefore, we employed metabolomics technology, 16S rRNA sequencing, and FMT methods to explore the interaction between DS and the gut microbiota in the prevention and treatment of ALI.

To ensure the standardization of DS medicinal herbs, this study employed UPLC-Q-TOF-MS technology for qualitative analysis, identifying a total of 37 chemical components in DS. Components such as Kaempferol, Quercetin-3-O-glucoside, and Aurantiamide present in DS have been widely studied due to their anti-inflammatory properties. Previous research has confirmed that Orientin, a flavonoid component with antioxidant capabilities, is considered a promising nutritional health product for patients with liver injury^33^. Similarly, Aurantiamide can exert an inhibitory effect on LPS-induced neuroinflammation by suppressing the NF-κB, JNK, and p38 MAPK pathways^34^. Loliolide can significantly accelerate LXR protein degradation by enhancing ubiquitination, thereby inhibiting adipogenesis. These findings suggest that by enhancing LXR ubiquitination to reduce adipogenesis, Loliolide and Pinoresinol may become potential complementary therapeutic candidates for non-alcoholic fatty liver disease^35^.

In our study, we utilized CCL_4_ to establish an ALI model. CCL_4_, as a compound capable of inducing hepatocyte necrosis, can partially recreate human liver disease patterns in animals, and thus has long been used by researchers as a modeling agent to induce various ALI models ^36^. During our experimentation, we observed early symptoms of liver fibrosis in our model, such as extensive fatty degeneration and necrotic fibrous septa. We conducted routine rat experiments, antibiotic treatments, and FMT experiments. Ultimately, we demonstrated that DS significantly alleviated ALI, improved liver function, and promoted the recovery of intestinal barrier function in a microbiota-dependent manner, thereby effectively inhibiting the progression of ALI.

When liver tissue is invaded by CCL_4_, the levels of AST, ALT, and AKP increase significantly ^37–39^. The expression changes of α-SMA and collagen-Ⅰ are interrelated, jointly promoting the development of ALI. High expression of α-SMA indicates the activation of hepatic stellate cells, which secrete large amounts of collagen-Ⅰ, leading to excessive deposition of extracellular matrix ^40^. Our research findings demonstrate abnormal liver function in MOD rats, with notably elevated expressions of α-SMA and collagen-Ⅰ, compromised intestinal barrier, and decreased expressions of ZO-1 and Claudin. Both the DSH and AMF groups exhibited reduced expressions of α-SMA and collagen-Ⅰ in liver tissue and increased expressions of ZO-1 and Claudin in colonic tissue, emphasizing their beneficial effects on liver tissue and barrier function. These discoveries underscore DS’s ability to inhibit ALI and promote the restoration of intestinal barrier function in a gut microbiota-dependent manner.

Considering the crucial role played by gut microbiota in the mechanism of ALI, we analyzed the impact of DS on gut microbiota and the influence of gut microbiota on DS-mediated ALI suppression. Our research findings indicate that DS can counteract the ecological imbalance caused by HI in the gut. It is noteworthy that DS increased the abundance of symbiotic probiotics such as *Lactobacillus*, *Bacilli*, *HT002*. *Lactobacillus* plays a significant role in protecting against liver injury, primarily through mechanisms such as regulating intestinal flora balance, improving intestinal barrier function, modulating metabolic pathways, and inhibiting inflammatory responses^41^.

More importantly, In FMT, it was discovered that DS improves ALI dependent microbiota alterations by enhancing fibrotic indicators and intestinal barrier function. Additionally, DS exhibited similar results to AMF in terms of gut microbiota modulation, with particular emphasis on the enrichment of *Lactobacillus*. Recent studies have shown that manipulating beneficial bacteria can restore intestinal barrier function, balance immune responses, alleviate intestinal inflammation, and even mitigate ALI^42^. The prominent presence of Lactobacillus suggests that DS may improve ALI effects. *Lactobacillus* has emerged as a potentially clinically useful probiotic, especially in the prevention and treatment of conditions like ALI, liver fibrosis^43^, and enteritis^44^. Meanwhile, studies have confirmed that *Lactobacillus* intervention can enhance the Th1 immune response and attenuate the Th2 immune response in ovalbumin-sensitized splenic lymphocytes^45^. In both the CON and AMF groups, *Firmicutes*, as the dominant bacterial group, can degrade dietary fiber to produce metabolites beneficial to health. These metabolites, which may include short-chain fatty acids, can modulate intestinal immune function and maintain the integrity of the intestinal barrier, thereby promoting intestinal health^46^. Furthermore, we discovered the emergence of the primary bile acid biosynthesis pathway in our non-targeted metabolomics analysis, which has significant implications for the bacterial group *Romboutsia*. Bile acids are crucial metabolites of the liver and play a vital role in regulating the structure and function of gut microbiota. Simultaneously, gut microbiota can also affect liver health by modulating bile acid metabolism. Research suggests that *Romboutsia* may be involved in the metabolic process of bile acids, influencing their composition and concentration, and thereby impacting liver health^47^. We plan to investigate these functions in future studies. Overall, DS improves ALI by increasing the abundance of probiotics, particularly the enriched *Lactobacillus* flora..

Tryptophan metabolism plays a pivotal role in various diseases, as its metabolites can influence the body’s immune response and inflammatory status through multiple pathways. Studies have indicated that modulation of tryptophan metabolism can activate the AhR signaling pathway, thereby alleviating inflammatory responses to a certain extent^48^. Furthermore, the inhibition of glycerophospholipid metabolism is closely associated with the regulation of inflammatory reactions. By regulating this metabolic pathway, it is possible to affect the stability of cell membranes and signal transduction, thereby influencing inflammatory responses ^49^. *Lactobacillus*, as a probiotic, has a close correlation between its abundance and the balance of gut microbiota. Research has found that an increase in *Lactobacillus* can influence tryptophan metabolism and glycerophospholipid metabolism by modulating the composition of gut microbiota, thereby providing some relief from ALI ^50^. By regulating these metabolic pathways, *Lactobacillus* can enhance the body’s antioxidant capacity and reduce the expression of inflammatory factors, exerting a protective effect on ALI. Joint analysis of metabolomics and gut microbiota has revealed a positive correlation between *Lactobacillus* and tryptophan metabolites such as Indoleacetaldehyde and N-Methyltryptamine. Conversely, there is a negative correlation with glycerophospholipid metabolic components such as PC(22:6(4Z,7Z,10Z,13Z,16Z,19Z)/18:1(9Z)), PC(22:5(7Z,10Z,13Z,16Z,19Z)/18:1(9Z)), PC(22:5(7Z,10Z,13Z,16Z,19Z)/16:0), and PC(20:1(11Z)/16:1(9Z)). We hypothesize that *Lactobacillus* improves ALI by activating the tryptophan metabolic pathway and inhibiting the glycerophospholipid pathway

Increasing evidence suggests that gut microbiota-derived metabolites serve as key signaling molecules in host-microbe crosstalk. Representative studies have shown that pectin supplementation alters the gut microbiota of mice, increases microbial tryptophan metabolites, and reduces alcohol-induced HI by activating the AhR pathway ^51, 52^. Our experimental results demonstrate that DS treatment significantly alleviates CCL_4_-induced ALI in rats by upregulating the expression of AhR and its downstream effector molecule Cyp1a1 in liver tissue, thereby activating the AhR signaling pathway. In our study, fecal metabonomics data indicated that DS modulates tryptophan metabolic pathways and upregulates beneficial secondary tryptophans such as L-Kynurenine and Indole-3-carboxaldehyde. Correlation analysis further revealed an association between Indole-3-carboxaldehyde and Lactobacillus abundance. Previous research suggests that the combination of Indole-3-carboxaldehyde with anti-fibrotic drugs may more effectively inhibit the progression of liver fibrosis, and its use with immunomodulators may better regulate intestinal mucosal immune homeostasis, further reducing liver inflammation ^53^. In summary, these findings indicate that DS regulates tryptophan metabolism, particularly the beneficial modulation of Indole-3-carboxaldehyde, in a gut microbiota-dependent manner, thereby inhibiting ALI.

## 5 Conclusion

In summary, our findings suggest that DS exerts inhibitory effects on ALI through the modulation of tryptophan metabolism mediated by gut microbiota. It effectively promotes the repair of intestinal barrier damage and reduces the expression of inflammatory factors, thereby suppressing ALI. Our study highlights the potential of DS as a modulator of gut microbiota, exerting inhibitory effects on ALI in a gut microbiota-dependent manner. These discoveries provide novel insights into the development of new natural medicinal agents for the prevention and treatment of ALI. However, since the content of individual components in DS extract was not accurately determined in this study, it remains uncertain whether variations in the composition of the extract could influence our results. Therefore, the relationship between different compositions of DS extract and their hepatoprotective effects, as well as the optimization of extraction methods to obtain DS extracts with optimal hepatoprotective properties, will be the focus of our future investigat.

## Ethics statement

The animal experiments were approved by the Ethical Approval Document of Baotou Medical College: BYDLR [2023] No. 38)

## Data Availability Statement

The datasets used and/or analyzed during the current study are available from the corresponding author upon reasonable request.

## CRediT authorship contribution statement

**Xiaolei Jiang**: Formal analysis, Investigation, Visualization, Writing – original draft. **Yafeng Zhuang**: Formal analysis, Visualization, Validation. **Tianwei Meng**: Investigation, Validation. **Jiali Liu**& **Tiancheng Meng**: Formal analysis. **Hongyu Meng**: Writing – review & editing. **Hong Chang**: Conceptualization, Project administration, Writing – review & editing.

## Abbreviations

ADS: Administration of *Dianthus superbus* L. to the pseudo-germ-free acute liver injury model
AhR: Aryl hydrocarbon receptor
ALT: Glutamate pyruvate transaminase
AMF: Administration of *Dianthus superbus* L. fecal microbiota solution to the pseudo-germ-free acute liver injury model
AMOD: Pseudo-germ-free acute liver injury model
AST: Glutamic-oxal(o)acetic transaminase
CCl_4_: Carbon tetrachloride
Collagen-Ⅰ: Collagen type I
Cyp1a1: Cytochrome P450 Family 1 Subfamily A Member 1
DS: *Dianthus superbus* L.
DSH: High-dose *Dianthus superbus* L.
DSL: Low-dose *Dianthus superbus* L.
DSM: Medium-dose *Dianthus superbus* L.
FMT: Fecal Microbiota Transplantation
OPLS-DA: Orthogonal Partial Least Squares-Discriminant Analysis PCA Principal Component Analysis
PcoA: Principal Coordinates Analysis
PLS-DA: Partial Least Squares Discrimination Analysis
Sily: Silibinin
VIP: Variable Importance in Projection TNF-α Tumor Necrosis Factor-alpha
ZO-1: Zonula Occludens-1
α-SMA: α-smooth Muscle Actin

## References

[1] Ghabril M, Vuppalanchi R, Chalasani N. Drug-Induced Liver Injury in Patients With Chronic Liver Disease.Liver Int 45:e70019 (2025).

[2] Morris SM, Chauhan A. The role of platelet mediated thromboinflammation in acute liver injury.Front Immunol 13:1037645 (2022).

[3] Yuan X, Zhou Y, Sun J, Wang S, Hu X, Li J, et al. Preventing acute liver injury via hepatocyte-targeting nano-antioxidants.Cell Prolif 56:e13494 (2023).

[4] Oh HA, Kim YJ, Moon KS, Seo JW, Jung BH, Woo DH. Identification of integrative hepatotoxicity induced by lysosomal phospholipase A2 inhibition of cationic amphiphilic drugs via metabolomics.Biochem Biophys Res Commun 607:1–8 (2022).

[5] Yang H, Park T, Park D, Kang MG. Trovafloxacin drives inflammation-associated drug-induced adverse hepatic reaction by changing macrophage polarization.Toxicol In Vitro 82:105374 (2022).

[6] Jaeschke H, Ramachandran A. Does acetaminophen hepatotoxicity involve apoptosis, inflammatory liver injury, and lipid peroxidation?J Biochem Mol Toxicol 35:e22718 (2021).

[7] Chen T, Shuang R, Gao T, Ai L, Diao J, Yuan X, et al. OPA1 Mediated Fatty Acid β-Oxidation in Hepatocyte: The Novel Insight for Melatonin Attenuated Apoptosis in Concanavalin A Induced Acute Liver Injury.J Pineal Res 76:e70010 (2024).

[8] Zhang SB, Qian J, Miao H, Zhang S, Hu Y, Liu P, Xu L. Mulberroside A ameliorates CCl4-induced liver fibrosis in mice via inhibiting pro-inflammatory response.Food Science & Nutrition (2023).

[9] Li H, Ye F, Li Z, Peng X, Wu L, Liu Q. The response of gut microbiota to arsenic metabolism is involved in arsenic-induced liver injury, which is influenced by the interaction between arsenic and methionine synthase.Environment International (2024).

[10] Huang H, Fang F, Jia Z, Peng W, Wu Y. Influences of Oral Administration of Probiotics on Posthepatectomy Recovery in Patients in Child-Pugh Grade.Computational and Mathematical Methods in Medicine (2022).

[11] Shang H, Huang C, Xiao Z, Yang P, Zhang S, Hou X, Zhang L. Gut microbiota-derived tryptophan metabolites alleviate liver injury via AhR/Nrf2 activation in pyrrolizidine alkaloids-induced sinusoidal obstruction syndrome.Cell and Bioscience (2023).

[12] Xie X, Zhang L, Yuan S, Li H, Zheng C, Xie S, et al. Val-Val-Tyr-Pro protects against non-alcoholic steatohepatitis in mice by modulating the gut microbiota and gut-liver axis activation.Journal of Cellular and Molecular Medicine (2021).

[13] Liu X, Zhao K, Yang X, Zhao Y. Gut Microbiota and Metabolome Response of Decaisnea insignis Seed Oil on Metabolism Disorder Induced by Excess Alcohol Consumption.Journal of Agricultural and Food Chemistry (2019).

[14] Bai X, Duan Z, Deng J, Zhang Z, Fu R, Zhu C, Fan D. Ginsenoside Rh4 inhibits colorectal cancer via the modulation of gut microbiota-mediated bile acid metabolism.Journal of Advanced Research (2024).

[15] Peng W, Meng D, Yue T, Wang Z, Gao Z. Effect of the apple cultivar on cloudy apple juice fermented by a mixture of Lactobacillus acidophilus, Lactobacillus plantarum, and Lactobacillus fermentum.Food Chemistry (2020).

[16] Liu C, Zheng J, Ou X, Han Y. Anti-cancer Substances and Safety of Lactic Acid Bacteria in Clinical Treatment.Frontiers in Microbiology (2021).

[17] Fang TJ, Guo JT, Lin MK, Lee MS, Chen YL, Lin WH. Protective effects of Lactobacillus plantarum against chronic alcohol-induced liver injury in the murine model.Applied Microbiology and Biotechnology (2019).

[18] Li S. Modulation of immunity by tryptophan microbial metabolites.Frontiers in Nutrition (2023).

[19] Jiang Z-M, Zeng S-L, Huang T-Q, Lin Y, Wang F-F, Gao X-J, et al. Sinomenine ameliorates rheumatoid arthritis by modulating tryptophan metabolism and activating aryl hydrocarbon receptor via gut microbiota regulation.Science Bulletin (2023).

[20] Chen C, Cao Z, Lei H, Zhang C, Wu M, Huang S, et al. Microbial Tryptophan Metabolites Ameliorate Ovariectomy-Induced Bone Loss by Repairing Intestinal AhR-Mediated Gut-Bone Signaling Pathway.Advanced Science (2024).

[21] Zhao Y, Li B, Deng H, Zhang C, Wang Y, Chen L, Teng H. Galangin Alleviates Alcohol-Provoked Liver Injury Associated with Gut Microbiota Disorder and Intestinal Barrier Dysfunction in Mice.Journal of Agricultural and Food Chemistry (2024).

[22] Zhao J-H, Li J, Zhang X-Y, Shi S, Wang L, Yuan M-L, et al. Confusoside from Anneslea fragrans Alleviates Acetaminophen-Induced Liver Injury in HepG2 via PI3K-CASP3 Signaling Pathway.Molecules (2023).

[23] Chen Y, Wang Y, Jiang S, Xu J, Wang B, Sun X, Zhang Y. Red-fleshed apple flavonoid extract alleviates CCl4-induced liver injury in mice.Frontiers in Nutrition (2023).

[24] Xiao H, Zhu X. H., Ji MY, Yi l, Li M. H. A survey on the application of “bashaga”-type drugs in compound preparations in Inner Mongolia.Chinese Journal of Experimental Formulary;26:190–98(2020).

[25] Arujah. Study on the protective effect and mechanism of Digida-4-flavored soup on liver ischemia-reperfusion injury in rats [bachelor’s degree]. 2024.

[26] Ge H, Wang A. Effects of Qing Liver Jiuwei San on Raf/MEK/ERK signaling pathway in rats with liver fibrosis.Chinese Journal of Integrative Medicine;41:848–54(2021).

[27] Wang W, Zhou X, Zhang T, Shen Q, Xiong Q, Sun R. Progress in the study of chemical composition and biological activity of Dianthus spp.Anhui Agricultural Science;51:23–29(2023)

[28] Yuan Y, Che L, Qi C, Meng Z. Protective effects of polysaccharides on hepatic injury: A review.International Journal of Biological Macromolecules (2019).

[29] Zhang J. Study on the anti rheumatoid and antitumor active ingredients of Coldwater Seven and DS. Paper presented at.

[30] Zhi Y, Song Y, Zhang M, Wang J, Zhao B. Pharmacological effects of carbon tetrachloride-induced acute liver injury in mice.Chinese Journal of Traditional Chinese Medicine;40:107-09+275(2022).

[31] Han L, Huang A, Chen J, Teng G, Sun Y, Chang B, et al. Clinical characteristics and prognosis of non-APAP drug-induced acute liver failure: a large multicenter cohort study.Hepatology International (2023).

[32] Jiang H, Yu Y, Hu X, Du B, Shao Y, Wang F, et al. The fecal microbiota of patients with primary biliary cholangitis (PBC) causes PBC-like liver lesions in mice and exacerbates liver damage in a mouse model of PBC.Gut Microbes (2024).

[33] He X, Chen J, Mu Y, Zhang H, Chen G, Liu P, Liu W. The effects of inhibiting the activation of hepatic stellate cells by lignan components from the fruits of Schisandra chinensis and the mechanism of schisanhenol.Journal of Natural Medicines (2020).

[34] Yoon C-S, Kim D-C, Lee D-S, Kim K-S, Ko W, Sohn JH, et al. Anti-neuroinflammatory effect of aurantiamide acetate from the marine fungus Aspergillus sp. SF-5921: inhibition of NF-κB and MAPK pathways in lipopolysaccharide-induced mouse BV2 microglial cells.International Immunopharmacology (2014).

[35] Kim SY, Lee JY, Jhin C, Shin JM, Kim M, Ahn HR, et al. Reduction of Hepatic Lipogenesis by Loliolide and Pinoresinol from Lysimachia vulgaris via Degrading Liver X Receptors.Journal of Agricultural and Food Chemistry (2019).

[36] Tao W, Hua J, Guocai L, Bojun Y, Jiahong S. New developments in carbon tetrachloride hepatotoxicity studies.Journal of Toxicology:324–27(2008).

[37] Gong Q, Zeng Z, Jiang T, Bai X, Pu C, Hao Y, Guo Y. Anti-fibrotic effect of extracellular vesicles derived from tea leaves in hepatic stellate cells and liver fibrosis mice.Frontiers in Nutrition (2022).

[38] Song J, Ren L, Ren Z, Ren X, Qi Y, Qin Y, et al. SIRT1-dependent mitochondrial biogenesis supports therapeutic effects of 4-butyl-polyhydroxybenzophenone compounds against NAFLD.European Journal of Medicinal Chemistry (2023).

[39] Wang P, Li J, Ji M, Pan J, Cao Y, Kong Y, et al. Vitamin D receptor attenuates carbon tetrachloride-induced liver fibrosis via downregulation of YAP.Journal of Hazardous Materials (2024).

[40] Ma M, Bao T, Li J, Cao L, Yu B, Hu J, et al. Cryptotanshinone affects HFL-1 cells proliferation by inhibiting cytokines secretion in RAW264.7 cells and ameliorates inflammation and fibrosis in newborn rats with hyperoxia induced lung injury.Frontiers in Pharmacology (2023).

[41] Ma P, Peng Y, Zhao L, Liu F, Li X. Differential effect of polysaccharide and nonpolysaccharide components in Sijunzi decoction on spleen deficiency syndrome and their mechanisms.Phytomedicine (2021).

[42] Ye J, Zhang C, Fan Q, Lin X, Wang Y, Azzam M, et al. Antrodia cinnamomea polysaccharide improves liver antioxidant, anti-inflammatory capacity, and cecal flora structure of slow-growing broiler breeds challenged with lipopolysaccharide.Frontiers in Veterinary Science (2022).

[43] Xiang H, Liu Z, Xiang H, Xiang D, Xiao S, Xiao J, et al. Dynamics of the gut-liver axis in rats with varying fibrosis severity.International Journal of Biological Sciences (2022).

[44] Lin W-S, Chueh T-L, Nagabhushanam K, Ho C-T, Pan M-H. Piceatannol and 3′-Hydroxypterostilbene Alleviate Inflammatory Bowel Disease by Maintaining Intestinal Epithelial Integrity and Regulating Gut Microbiota in Mice.Journal of Agricultural and Food Chemistry (2023).

[45] Yun X, Shang Y, Li M. Effect of Lactobacillus salivarius on Th1/Th2 cytokines and the number of spleen CD4⁺ CD25⁺ Foxp3⁺ Treg in asthma Balb/c mouse.International journal of clinical and experimental pathology (2015).

[46] Wang Q, Zhan X, Wang B, Wang F, Zhou Y, Xu S, et al. Modified Montmorillonite Improved Growth Performance of Broilers by Modulating Intestinal Microbiota and Enhancing Intestinal Barriers, Anti-Inflammatory Response, and Antioxidative Capacity.Antioxidants (2022).

[47] Cao Y-J, Huang Z-R, You S-Z, Guo W-L, Zhang F, Liu B, et al. The Protective Effects of Ganoderic Acids from Ganoderma lucidum Fruiting Body on Alcoholic Liver Injury and Intestinal Microflora Disturbance in Mice with Excessive Alcohol Intake.Foods (2022).

[48] Liu C, Wang Y, Sheng L, Zhang Y, Luo G, Ruan XZ, et al. 3-Hydroxypropionaldehyde modulates tryptophan metabolism to activate AhR signaling and alleviate ethanol-induced liver injury.Phytomedicine;139:156445. eng(2025).

[49] Cao Y, Fan X, Zang T, Li Y, Tu Y, Wei Y, et al. Gut microbiota causes depressive phenotype by modulating glycerophospholipid and sphingolipid metabolism via the gut-brain axis.Psychiatry Res;346:116392. eng(2025).

[50] Wang L, Xi M, Cao W, Qin H, Qin D, Chen S, et al. Electroacupuncture alleviates functional constipation by upregulating host-derived miR-205-5p to modulate gut microbiota and tryptophan metabolism.Front Microbiol;16:1517018. eng(2025).

[51] Wrzosek L, Ciocan D, Hugot C, Spatz M, Dupeux M, Houron C, et al. Microbiota tryptophan metabolism induces aryl hydrocarbon receptor activation and improves alcohol-induced liver injury.Gut (2020).

[52] Schanz O, Chijiiwa R, Cengiz SC, Majlesain Y, Weighardt H, Takeyama H, Förster I. Dietary AhR Ligands Regulate AhRR Expression in Intestinal Immune Cells and Intestinal Microbiota Composition.International Journal of Molecular Sciences (2020).

[53] D’Onofrio F, Renga G, Puccetti M, Pariano M, Bellet MM, Santarelli I, et al. Indole-3-Carboxaldehyde Restores Gut Mucosal Integrity and Protects from Liver Fibrosis in Murine Sclerosing Cholangitis.Cells (2021).

